# GADL1 is a multifunctional decarboxylase with tissue-specific roles in β-alanine and carnosine production

**DOI:** 10.1101/2020.02.18.954438

**Authors:** Elaheh Mahootchi, Selina Cannon Homaei, Rune Kleppe, Ingeborg Winge, Tor-Arne Hegvik, Roberto Megias-Perez, Christian Totland, Floriana Mogavero, Anne Baumann, Jeffrey Glennon, Hrvoje Miletic, Petri Kursula, Jan Haavik

**Author notes:** **Corresponding Author:** Professor Jan Haavik, Department of Biomedicine, University of Bergen, Jonas Lies vei 91, 5020 Bergen, Norway.

## Abstract

Carnosine and related β-alanine-containing peptides are believed to be important antioxidants, pH-buffers and neuromodulators. However, their biosynthetic routes and therapeutic potential are still being debated. This study describes the first animal model lacking the enzyme glutamic acid decarboxylase-like 1 (GADL1). We show that Gadl1^-/-^mice are deficient in β-alanine, carnosine and anserine, particularly in the olfactory bulb, cerebral cortex, and skeletal muscle. Gadl1^-/-^mice also exhibited decreased anxiety, increased levels of oxidative stress markers, alterations in energy and lipid metabolism, and age-related changes. Examination of the GADL1 active site indicated that the enzyme may have multiple physiological substrates, including aspartate and cysteine sulfinic acid, compatible with organ-specific functions. Human genetic studies show strong associations of the GADL1 locus with plasma levels of carnosine, subjective well-being, and muscle strength, also indicating a role for β-alanine and its peptide derivatives in these traits. Together, this shows the multifaceted and organ specific roles of carnosine peptides and establishes Gadl1 knockout mice as a versatile model to explore carnosine biology and its therapeutic potential.

## Introduction

Carnosine (β-alanyl-L-histidine) is one of several dipeptides of β-alanine and histidine that are found in high concentrations in vertebrate tissues, particularly in brain and skeletal muscle (SKM) tissues. In humans, carnosine and acetylcarnosine are most abundant. In contrast, many animals mainly synthesize the related β-alanine-containing peptides anserine or ophidine/balenine through methylation of the imidazole moiety of carnosine in the 1-or 3-position(1–3). Here, we refer to this family of related peptides as “carnosine peptides”.

Carnosine peptides may have multiple biological functions, including calcium regulation, pH buffering, metal chelation, and antioxidant effects(2). β-alanine, as well as its dipeptide derivatives, may also be neurotransmitters or neuromodulators in the central nervous system (CNS), especially in OB. In mammals, including mice, only two tissues have carnosine concentrations in the millimolar range, *i.e.* SKM and olfactory bulb (OB)(2, 4). Various disorders and dysfunctions have been linked to alterations in β-alanine and carnosine metabolism. In β-alaninemia and carnosinemia, decreased degradation of these compounds is associated with neurological symptoms(5, 6). Conversely, it has been suggested that some age-related and neurological diseases, e.g. Alzheimer’s disease, Parkinson’s disease, multiple sclerosis (MS) and cancer, as well as diabetes complications, are related to carnosine deficiency(7).

β-alanine supplementation increases carnosine content and improves contractility and muscle performance of human and rodent SKM, especially during short exercise intervals(8, 9). Such findings have led to the widespread use of dietary β-alanine supplements, particularly among athletes and soldiers(10). β-alanine and carnosine dietary supplementation have also shown some promising effects in the treatment of depression, anxiety and autism symptoms in humans(11) and animal models(12, 13).

As most carnosine peptides are hydrolyzed after oral ingestion, either in enterocytes or in the blood circulation, the body mainly depends on *de novo* synthesis of these peptides. Dietary supplementation has side effects and is ineffective since a daily intake of several grams is needed to significantly increase the muscle β-alanine and carnosine content(14). The lack of suitable model systems and the need to supply large pharmacological quantities of carnosine peptides has caused much uncertainty regarding the mechanisms of action and safety of these compounds. A more stable, synthetic analog of carnosine, carnosinol, was recently shown to protect against metabolic dysregulation and oxidative damage(15).

Synthesis of β-alanine appears to be a rate-limiting factor for carnosine peptide levels in mammals(16). In a large genome-wide association (GWA) study on plasma metabolites, it was estimated that 86% of the variation in carnosine levels could be attributed to genetic factors, making this peptide the most heritable metabolite among all compounds examined(17). Still, uncertainty remains regarding the identity of the genes and proteins involved in its synthesis. Carnosine synthase 1 (encoded by *CARNS1*) produces carnosine from β-alanine and histidine and homocarnosine from γ-aminobutyric acid (GABA) and histidine. In animal tissues β-alanine may be produced by reductive degradation of uracil(18). In contrast, some bacteria produce β-alanine by α-decarboxylation of aspartic acid (Asp), catalyzed by aspartate decarboxylase with the aid of a covalently bound pyruvoyl cofactor(19). While in insects, an analogous vitamin B6 (pyridoxal phosphate, PLP)-dependent enzyme *L*-aspartate-α-decarboxylase has evolved by convergent evolution(20, 21). Intriguingly, the formation of carnosine in mouse SKM is dependent on vitamin B6, implicating a PLP-dependent enzyme in its synthesis also in animals(22).

Based on its sequence similarity to glutamic acid decarboxylase, the PLP-dependent enzyme glutamic acid decarboxylase-like protein 1 (GADL1; acidic amino acid decarboxylase) has been assigned a possible role in GABA synthesis(21). Partially reflecting this connection, *GADL1* has been suggested to be associated with lithium response in bipolar patients(23). Although this genetic association was not replicated in other clinical samples(24), these findings have triggered interest in the *in vivo* investigation of GADL1 function and biochemical properties. Cysteine sulfinic acid (CSA) is one of the substrates of GADL1 and it was suggested that GADL1 is involved in the biosynthesis of hypotaurine and taurine(21). However, it has also been reported that purified human(21) and mouse(25) GADL1 can synthesize β-alanine from Asp, although at very low rates *in vitro.* This is consistent with recent crystallographic studies, showing that GADL1 shares structural features with both CSAD and Asp decarboxylase(26). Moreover, a human genetic association study demonstrated that single nucleotide polymorphisms (SNPs) in the *GADL1* intron are strongly associated with blood levels of N-acetylcarnosine(27) that, in turn, are strongly correlated with carnosine levels(3). Based on these observations, we hypothesized that GADL1 could be involved in β-alanine and carnosine production in mammalian tissues.

Here, we describe the first *Gadl1* knockout (KO) mouse model and demonstrate an organ-specific role of GADL1 in the biosynthesis of carnosine peptides and protection against oxidative stress, particularly in the OB and SKM. To understand the substrate specificity of GADL1, we compared the 3-dimensional (3D) structures of mouse GADL1 and related enzymes. Finally, we investigate the association between common genetic variants in enzymes and transporters involved in carnosine homeostasis with multiple human traits and diseases and present an initial behavioral characterization of the *Gadl1* KO mouse.

## RESULTS

### Mice lacking GADL1 show age-related changes

*Gadl1^-/-^*(null) mice were generated using *Cre*/loxP technology. Due to the proximity of the *Gadl1* gene to the *Tgfbr2* gene that encodes a receptor with possible effects on growth and survival, we applied a conservative knocking out strategy, where only *Gadl1 exon 7*, coding for part of the PLP-binding active site of GADL1 was deleted (Fig. 1a). *Gadl1^+/+^, Gadl1^+/-^* and *Gadl1^-/-^*mice were successfully generated in the expected Mendelian ratios by breeding from *Gadl1*^+/-^mice. We observed no obvious physical abnormalities, and overall behavior and initial growth curves were similar across all genotypes. However, after 30 weeks of age, compared to *Gadl1^+/+^* mice, female and male *Gadl1^-/-^*mice exhibited relative growth retardation, compatible with age-related and possible degenerative changes (Fig. 1b-c).

**Fig. 1.**
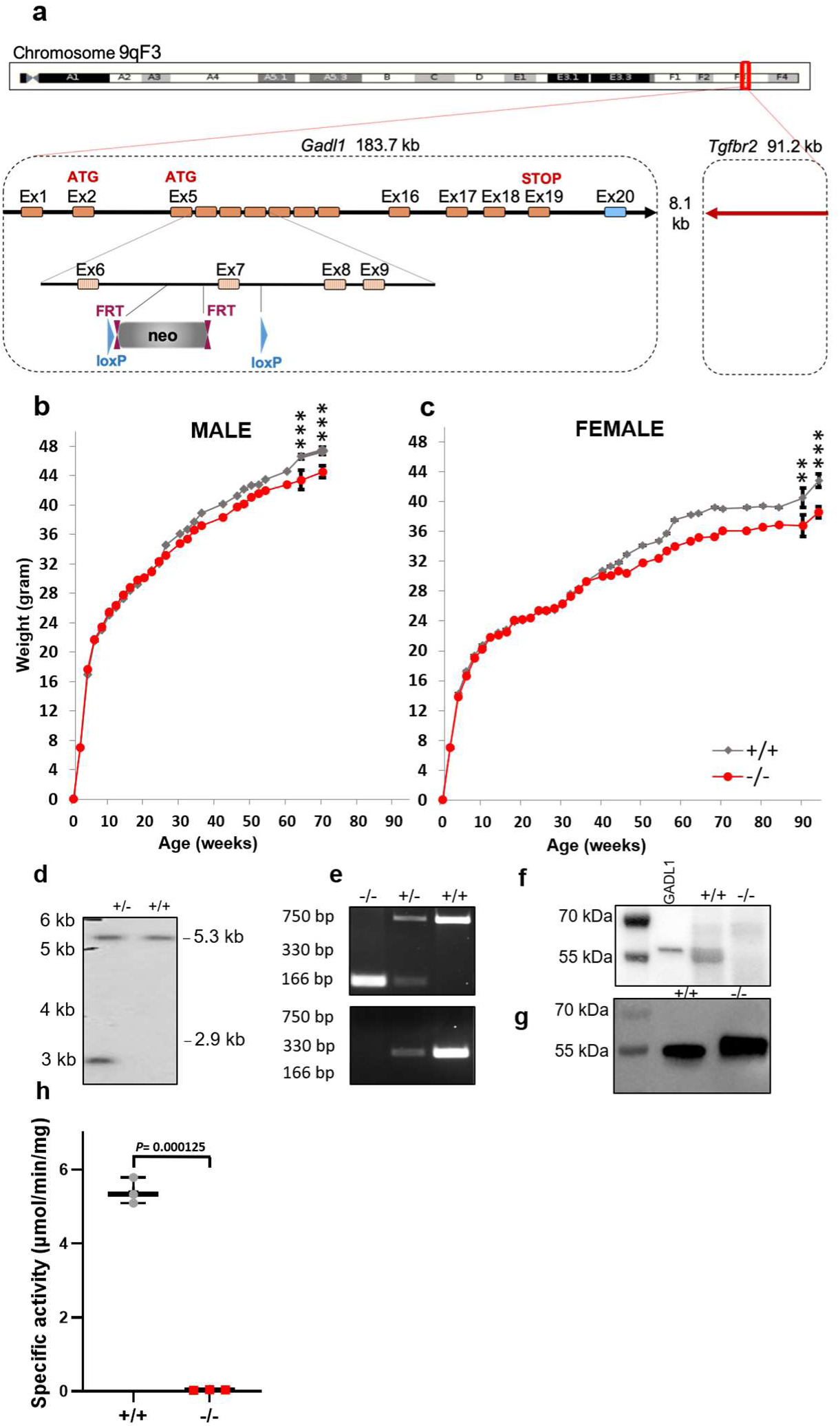
Generation of constitutive GADL1 KO mice. **a** Targeting strategy for knocking out exon 7 of the mouse *Gadl1* locus on chromosome 9. *Gadl1* coding sequences (hatched rectangles), non-coding exon portions (blue rectangles) and chromosome sequences (orange rectangles) are represented. The neomycin positive selection cassette is indicated between loxP sites (blue triangles) and FRT sites (plum triangles). **b, c** Growth curves of *Gadl1^+/+^* (n=4­34) and *Gadl1^-/-^*mice (n=4-40) from postnatal week 2 to 70 for **(b)** male and 90 weeks for **(c)** female. Presented as mean ± SD. Differences between genotypes were significant (*p*=0.0008 (64 weeks) and *p*=0.0005 (70 weeks) for males and *p*=0.0084 (90 weeks) and *p*=0.0001 (94 weeks) for females, respectively). **d** Southern blot analysis of genomic DNA from *Gadl1^+/+^* and *Gadl1^-/-^*. **e** Genotyping of the offspring from intercrosses of *Gadl1^+/-^*mice by PCR. The DNA band at 166 bp is the KO allele (primer 3), while bands at 330 bp (primer 2 and 3) and 750 bp (primer 1) are the WT alleles. **f** Representative western blots of OB samples from *Gadl1^+/+^* and *Gadl1^-/-^*mice (34 weeks old, female) using anti GADL1 antibody. Positive control was recombinant His-tagged GADL1 (2 ng, lane 1). **g** Western blot of recombinant His-tagged truncated *Gadl1^+/+^* and *Gadl1^-/-^*. **h** Enzyme activity towards CSA of recombinant His-tagged truncated *Gadl1^+/+^* and *Gadl1^-/-^*, *p* < 0.001.

Genomic DNA sequencing and Southern blot analyses confirmed the elimination of exon 7 (Fig. 1d-e). However, quantification of *Gadl1* mRNA from SKM and OB revealed that some *Gadl1* mRNA species could be detected across all genotypes. RNA sequencing showed that *Gadl1^-/-^*mice lacked *Gadl1* exons 7 and 8. This was confirmed using qRT-PCR of individual exons (see below). Bioinformatic analyses showed that the deletion of exons 7 and 8 was coupled to the generation of a new RNA splicing site in the mutated mice (Supplementary Fig. 1).

Western blotting confirmed that GADL1 protein was present in *Gadl1^+/+^* and *Gadl1^+/-^*, but not in *Gadl1^-/-^*mice. In OB extracted from *Gadl1^+/+^* mice, GADL1 appeared as a wide band, with an estimated molecular mass of 55-59 kDa (Fig. 1f). This corresponds to several predicted protein variants with 502 to 550 amino acids. Since exons 7 and 8 encode amino acids involved in cofactor binding, GADL1 lacking these amino acids was predicted to be enzymatically inactive. This was confirmed by expressing the protein lacking exons 7 and 8 in *E. coli* and comparing it to the full-length protein (Fig. 1g and Supplementary Fig. 2). The mutant protein yield was only 10% compared to that of the full-length protein. Furthermore, the mutant protein was completely devoid of enzyme activity (Fig. 1h). This demonstrated that the elimination of gene function was successful and that the *Gadl1^-/-^*mice did not have any residual GADL1 enzyme activity.

### Deletion of *Gadl1* perturbs carnosine metabolism

To explore the biological function(s) of GADL1, we performed untargeted LC-MS metabolomic analyses of eight different tissues from 20 *Gadl1^+/+^* mice and 21 *Gadl1*^-/-^ mice that were matched across genotype and sex (Fig. 2). The number of identifiable metabolites varied between 541-720 in the cerebral cortex, OB, SKM, liver, cerebellum, heart, serum, and kidney. Partial Least Square-Discriminant Analyses (PLS-DA) were used to evaluate metabolic differences between *Gadl1^+/+^* and *Gadl1*^-/-^mice. This allowed identification of suitable markers responsible for the metabolic differences by the projection by variable important projection (VIP) score.

**Fig. 2.**
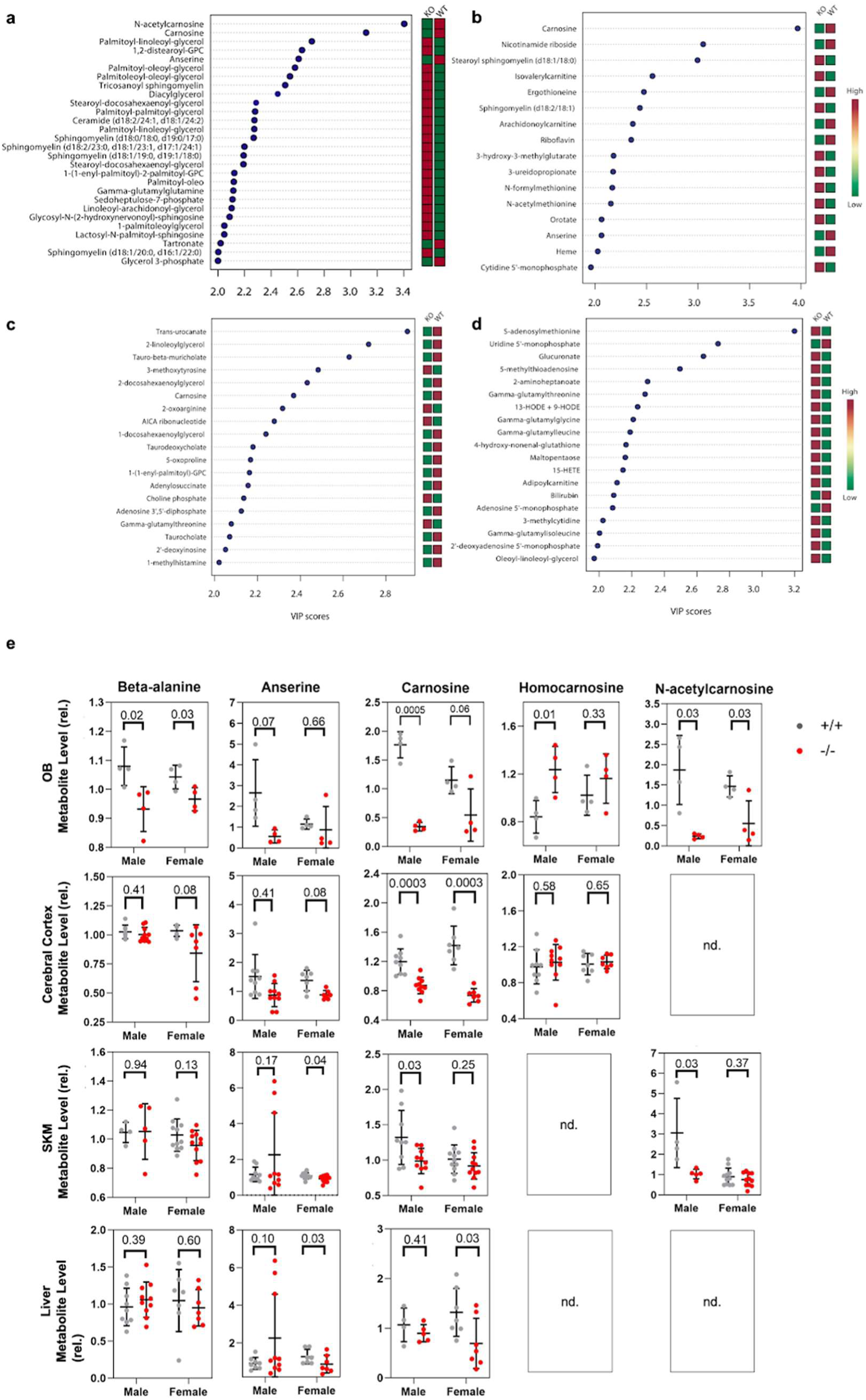
The deletion of *Gadl1* has tissue-specific effects on metabolite levels. **a-d** Top significant features of metabolites based on VIP scores > 2 of component 1 of PLS-DA. Untargeted metabolic profiling of **a** OB, **b** cerebral cortex, **c** SKM, and **d** liver tissue samples from *Gadl1^+/+^* (n=20) and *Gadl1^-/-^*(n=21) mice. **e** The relative levels of β­alanine and carnosine derivatives in *Gadl1^+/+^* (grey) and *Gadl1^-/-^*(red) mouse tissue.

The metabolic features with VIP scores higher than 2, responsible for the separation in each tissue are depicted in Fig. 2 (a-d). Carnosine peptides showed the strongest difference between the genotypes in the OB and the cerebral cortex (Fig. 2e). Carnosine was also among the most strongly affected metabolites in SKM, but not in the liver (Fig. 2e). Although GADL1 has been postulated to synthesize GABA from glutamate, the levels of GABA and glutamate in *Gadl1*^+/+^ and *Gadl1^-/-^*mice were similar in all tissues examined (fold difference 0.88-1.15 for glutamate and 0.96-1.53 for GABA, *p*>0.05). *Gadl1*^-/-^mice had significantly reduced levels of β-alanine in OB, but non-significant reduction in the cerebral cortex and SKM and unaltered levels in the liver. In contrast, carnosine, N-acetylcarnosine, and anserine were significantly depleted in all organs, except kidneys, cerebellum, and heart where this reduction was non-significant (data not shown). Together, this is consistent with the observed tissue distribution of GADL1 and demonstrates its key role in the metabolic pathway of β-alanine and carnosine peptides.

### High-resolution magic-angle spinning (MAS) NMR spectroscopy confirms the depletion of carnosine in *Gadl1^-/-^*mice

To determine the role of GADL1 on carnosine content in intact tissue, we used fresh OB tissue where we found the highest expression GADL1. We performed high-resolution magic-angle spinning ^1^H-NMR spectroscopy on OB from *Gadl1^+/+^* (n=4), *Gadl1^+/-^*(n=3) and *Gadl1^-/-^*(n=3) mice. Characteristic features of carnosine (Fig. 3a) could be identified in the aromatic regions of the spectrum, in accordance with quantification in tissue extracts (Fig. 3b). Compared to *Gadl1^+/+^*, *Gadl1^+/-^*and *Gadl1^-/-^*mice had 34% and 70% (*p* =0.0056) reduced carnosine content in the OB (Fig. 3c). To our knowledge, this is the first demonstration of carnosine measurement in intact brain tissue, also establishing MAS NMR spectroscopy as a fast and convenient way to determine carnosine in this tissue. There were no significant differences in the brain morphology of *Gadl1* genotypes (Fig. 3d-f).

**Fig. 3.**
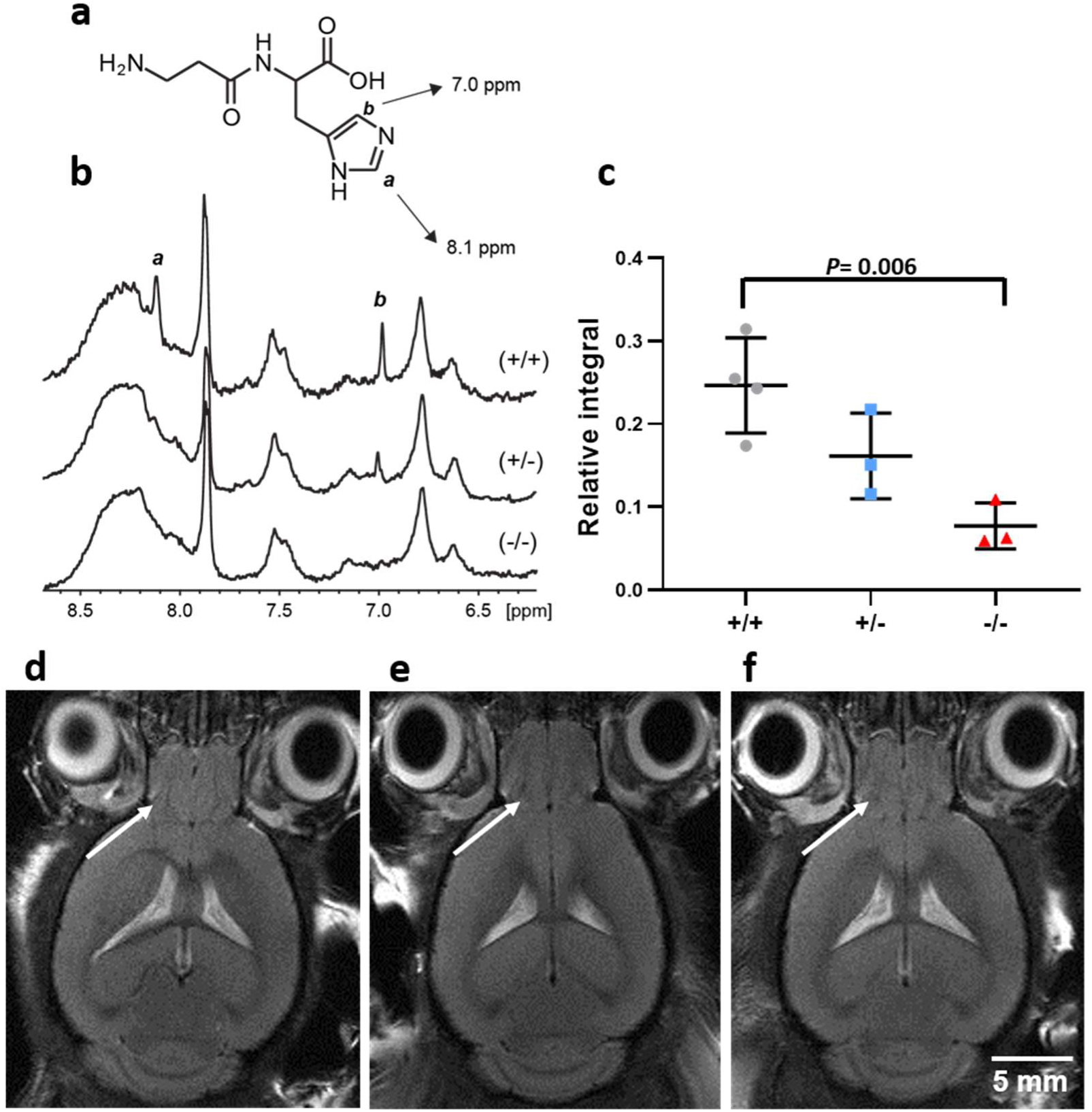
^1^H-NMR and MRI of mouse tissues. **a-c** Measurement of carnosine in intact OB tissue. **a** Chemical structure of carnosine. **b** MAS-^1^H-NMR spectra of OB tissue samples from *Gadl1^+/+^*, *Gadl1^+/-^*and *Gadl1^-/-^*male mice (12 weeks). The two hydrogens of the imidazole ring in carnosine are marked a and b. **c** Relative integral based on NMR results, presented as mean ± SD, **d-f** MRI of the brain in **d** *Gadl1^+/+^*, **e** *Gadl1^+/-^*and **f** *Gadl1^-/-^*mice. The arrow indicates the OB.

### Elimination of *Gadl1* alters the OB transcriptome

To determine the effects of GADL1 depletion on gene expression, we sequenced and compared OB mRNA levels from *Gadl1^+/+^, Gadl1^+/-^,* and *Gadl1^-/-^*mice (Fig. 4a-c). The top 25 up-and down-regulated genes (log2Fold change either less than −1 or greater than +1 and *p* ≤ 0.05) are shown in Supplementary Table 1 and Supplementary Fig.3. Pathway enrichment analysis showed that the strongest affected group of genes is involved in drug metabolism (KEGG pathway mmu00983, *p*=0.00016, q=0.014)(28) (Table 1). This pathway includes the *Upb1* gene (b-ureidopropionase 1), a transcript with a two-fold increase in *Gadl1^-/-^*mice (*p*=0.0154). UPB1 catalyzes the last step in the formation of β-alanine from pyrimidines (Fig. 7a). This is consistent with a compensatory mechanism for maintaining β-alanine synthesis in the absence of GADL1. Moreover, in OB of *Gadl1^-/-^*mice, we observed increased transcript levels of multiple isoforms of carboxylesterase 1 (Ces1), cytochrome P450 and myeloperoxidase (Fig. 4a-c, Table 1).

**Fig. 4.**
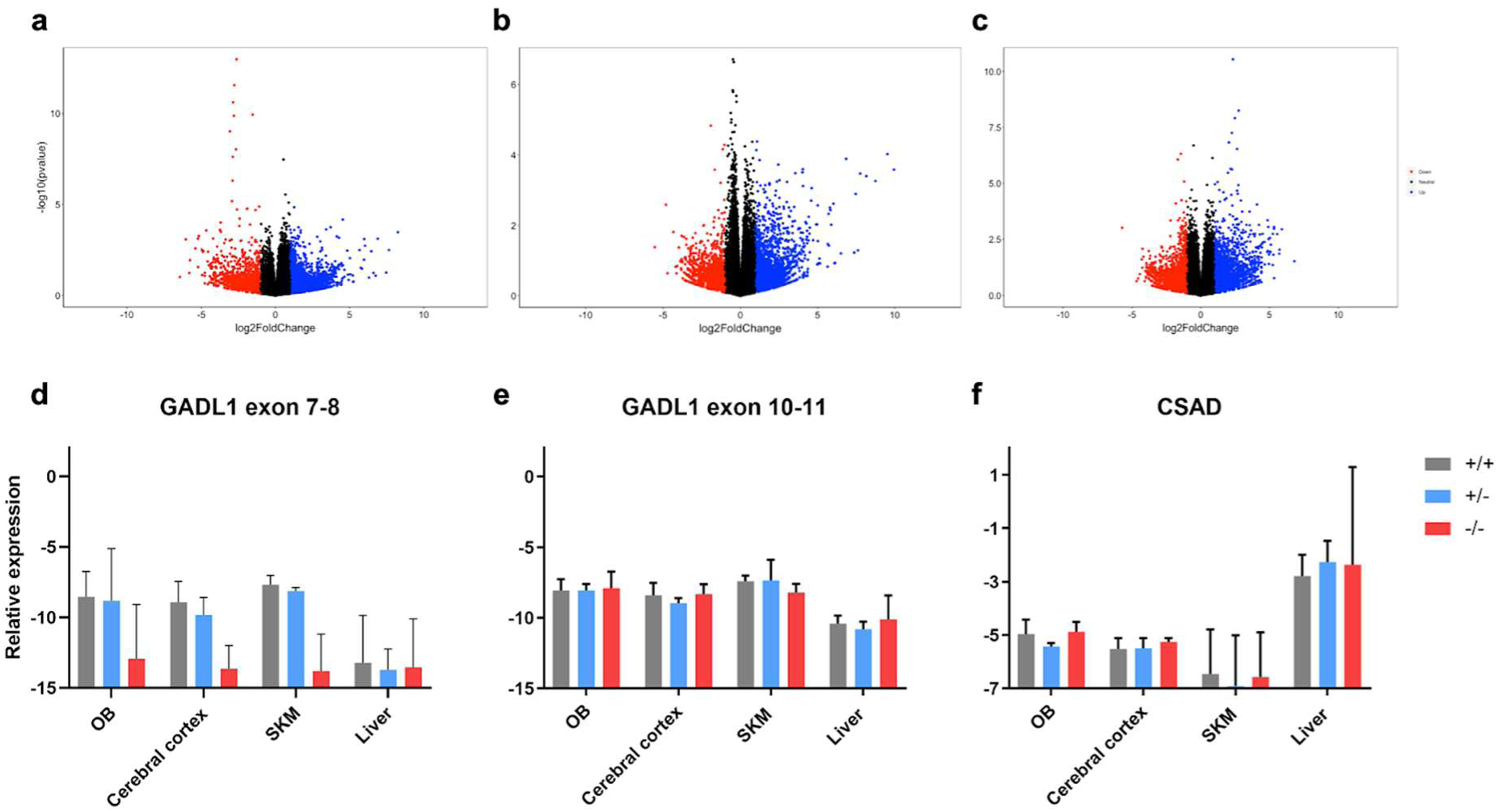
RNA sequence and qRT-PCR analysis of mouse tissues. **a-c** Relative RNA expression levels (volcano plots) in OB tissue. **a** *Gadl1^+/+^* to *Gadl1^+/-^*ratio, **b** *Gadl1^+/+^* to *Gadl1^-/-^*ratio, **c** *Gadl1^-/-^*to *Gadl1^+/-^*ratio. **d-f** qRT-PCR analysis of normalized mRNA expression in OB, brain, SKM, and liver tissues from 35 week old females *Gadl1^+/+^* (grey)*, Gadl1^+/-^*(blue) and *Gadl1^-/-^*(red) mice for **d** *Gadl1* exon 7 and 8**,e** *Gadl1* exon 10 and 11, and **f** CSAD. n=3 for each genotype. Presented on a Ln y-scale as mean of 2^ΔCt^ and upper limit (95%).

**Table 1.**
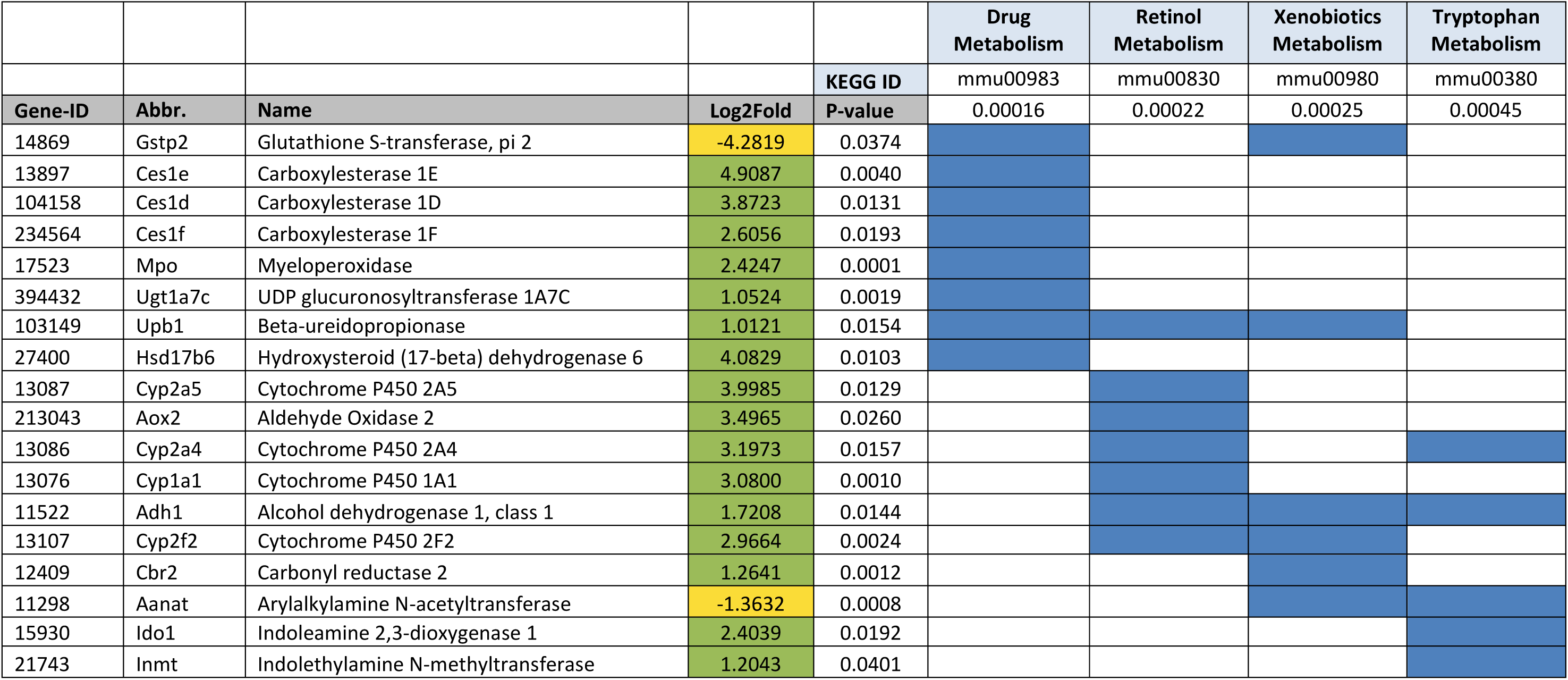
Pathway enrichment analysis comparing *Gadl1^-/-^*to *Gadl1^+/+^* (upregulated (green), downregulated (yellow)).

A gene ontology (GO) pathway analysis(28, 29) of up- or down-regulated genes (log2Fold change either less than −1 or greater than +1 and *p*≤0.05) showed significant alterations in 24 different biological processes, where the most significant alteration involves drug catabolism (GO:0042737, *p*=2.43e-07, q=0.0007). Furthermore, biological processes involving circadian rhythms and sleep cycles seemed to be altered. Dopamine receptors (*Drd1-3*) and adenosine A2a receptor (*Adora2a*) were the most consistently recurring genes throughout the GO analysis (Supplementary Table 2). It has been reported that the release of glutamate from terminals of carnosine containing olfactory neurons in OB glomeruli is modified by dopamine receptors(30). It is possible that the elimination of GADL1 and carnosine depletion also affected the expression of these modulatory transmitter receptors in the OB. Wu *et al.* recently reported that GADL1 overexpression inhibited KCTD12 expression(31). However, a comparison of *Gadl1^-/-^*and *Gadl1^+/+^* mice, showed that RNA levels of KCTD12 were unaffected by the deletion of *Gadl1* (*p*=0.62).

### Substrate specificity of GADL1

Many PLP-dependent enzymes have multiple substrates(32). To explore whether the divergent metabolic changes observed in *Gadl1*^-/-^mice could be secondary to changes in β­alanine levels or were due to the parallel chemical conversion of multiple substrates by GADL1, we solved the crystal structure of mouse GADL1(26). The structure of the closely related CSAD has been solved earlier, but not published (PDB entry 2JIS). With available crystal structures of both GADL1, CSAD, and GAD(33), it is possible to identify determinants of substrate specificity in these acidic amino acid decarboxylases. GADL1 and CSAD have different, although slightly overlapping, physiological functions(25). GADL1, through the conversion of Asp to β-alanine, is important in carnosine peptide biosynthesis, while CSAD catalyzes the major pathway in taurine synthesis, using CSA as substrate. Despite similar chemical structures of the two substrates (Fig. 5a), these two highly similar enzymes can distinguish between them. Both Asp and CSA are smaller than Glu (Fig. 5b), explaining the inability of GADL1 and CSAD to function as glutamate decarboxylases (Fig. 5c). The carboxyl group in Asp is planar, while the sulfinic acid moiety of CSA is tetrahedral and has slightly longer bonds; effectively, CSA is slightly larger and can perhaps more specifically accommodate to a binding site due to its shape and charge.

**Fig. 5.**
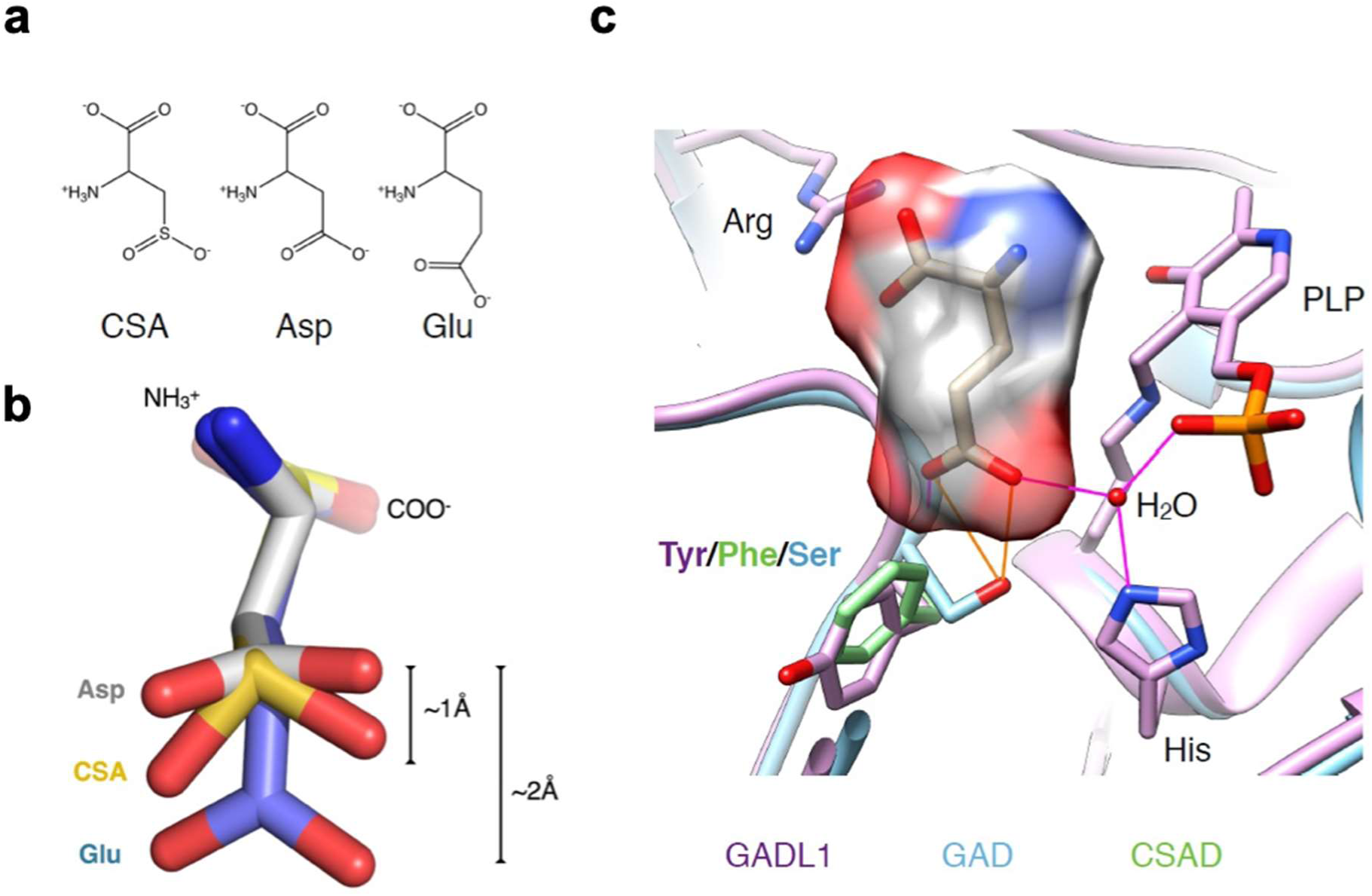
Substrate specificity. **a** Chemical structure comparison of GADL1 substrates CSA, Asp, and Glu. **b** 3D substrate structures to show the size and shape differences of Glu compared to CSA and Asp. **c** Active sites of GADL1, GAD, and CSAD and the predicted mode of binding of Glu to GAD. The prediction is based on the complex between GAD and the inhibitor chelidonic acid(34).

As the chemical reaction catalyzed by the enzymes is identical, we can assume that the reactive groups bind in an identical manner, and the differences lie in recognizing the side chain of the substrate. GAD uses Glu as a substrate, while GADL1 acts on Asp, indicating that side-chain length is one determinant of productive binding. For binding a negatively charged side chain in the specificity pocket, one expects to find positive charge potential, in the form of amino groups. Indeed, for GAD, such interactions can be conceived; two backbone NH groups as well as a water molecule coordinated by a conserved His residue coordinate the side-chain carboxyl group in the model of the substrate complex, together with a Ser side chain. How is GADL1 different? The major difference is the substitution of this Ser with a Tyr residue (Fig. 5c), effectively making the binding cavity smaller. Interactions with the peptide backbone and the His-water unit are likely conserved for Asp binding to GADL1. The specificity pocket contains a Tyr (GADL1) or Phe (CSAD) residue, and the slightly different conformations of these residues are enough to cause this difference in substrate specificity.

### Tissue-specific effects of GADL1 on the synthesis of taurine and its derivatives

Based on the ability of purified GADL1 to produce hypotaurine from CSA, it has been suggested that the primary biological function of GADL1 may be in taurine synthesis(21). However, the concentrations of the substrates and products in taurine synthesis were unaltered in the *Gadl1^-/-^*mice in the brain, OB and liver. In contrast, levels of taurine derivatives were moderately reduced in SKM in *Gadl1^-/-^*(Supplementary Fig. 4). This is consistent with the observations that taurine mainly is synthesized by other routes, such as by CSAD(35) and our previous finding that CSAD is expressed in low amounts in SKM(25). In all tissues examined, CSAD protein has been reported to be more abundant than GADL1, and purified CSAD has a 5-35 fold higher specificity constant for decarboxylation of CSA than GADL1(25). Moreover, as shown here, the qRT-PCR analysis showed that the levels of *Csad* mRNA were at least 10-fold higher than levels of *Gadl1* and were unaffected by the deletion of *Gadl1 exons* (Fig. 4f). Thus, we conclude that in organs with a high content of CSAD, GADL1 has a minor role in taurine synthesis. In the absence of CSAD, levels of taurine in mouse plasma, liver, and brain decreased by 70-90%, with the largest decrease in liver and the smallest in the brain(35). This remaining taurine biosynthetic capacity has been attributed to an alternative biosynthetic route of taurine catalyzed by cysteamine dioxygenase from cysteamine, generated from coenzyme A(36). However, our results indicate that GADL1 has a small taurine biosynthetic capacity *in vivo* that may contribute to the overall production of taurine.

### Oxidative stress markers

Several lines of evidence support a role of carnosine peptides in protection against oxidative stress(1, 2). To investigate the role of GADL1 in antioxidant defense, we compared levels of oxidative stress markers in *Gadl1^+/+^* and *Gadl1^-/-^*mice. *Gadl1^-/-^*mice had increased levels of oxidative stress markers, including methionine sulfoxide and γ-glutamyl peptides (Supplementary Fig. 5). These alterations are compatible with elevated glutathione synthesis and consistent with elevated oxidative stress. Interestingly, some oxidative stress markers were strongly increased even in tissues with a moderate decrease in carnosine peptides, such as the liver. We also compared tissue levels of the antioxidant enzymes SOD1 (CuZnSOD), SOD2 (MnSOD), and glutathione reductase (GSR) in *Gadl1^+/+^* and *Gadl1*^-/-^mice (OB, cerebral cortex, and SKM) using western blotting (Fig. 6). Surprisingly, the most obvious difference was detected in the OB where, the *Gadl1^-/-^* mice had a 3-fold increase in GSR levels (*p*=0.0145) (Fig. 6a).

**Fig. 6.**
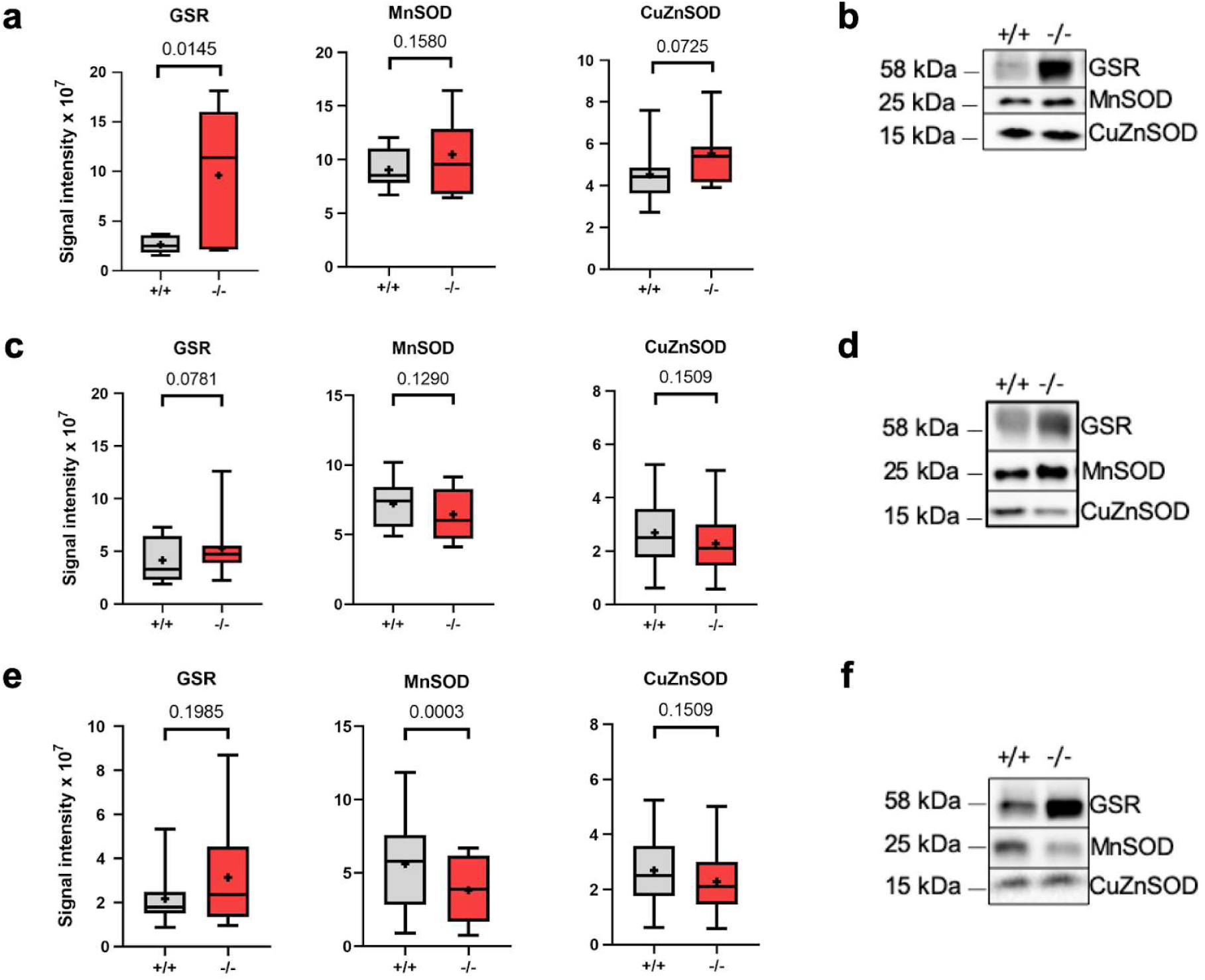
*Gadl1^-/-^*mice have altered levels of antioxidant enzymes. **a-e** Representative western blots **(b, d, f)** and normalized protein expression **(a, c, e)** for glutathione reductase (GSR), superoxide dismutase 1 (CuZnSOD), and SOD2 (Mn) in **(a, b)** OB, **(c, d)** cerebral cortex and **(e, f)** skeletal muscle tissue from female *Gadl1^+/+^* and *Gadl1^-/­^*mice.

*Gadl1^-/-^*mice had increased levels of many lipid species, including sphingolipids, with the strongest effects in OB (Figs. 2a-d, Supplementary Fig. 6). Notably, complex sphingolipids accumulate under inflammation and oxidative stress, conditions which *Gadl1^-/-^*mice seem more susceptible to. Moreover, sphingolipids may alter muscle contractility(37).

### Animal behavior and tissue morphology

To examine the effects of *Gadl1* deletion on brain function, we tested a range of mouse behaviors in *Gadl1^+/+^* (n=13) and *Gadl1^-/-^*(n=13) 22 week old male mice. In the open field, the latency to first enter the center was significantly different between genotypes, with *Gadl1^-/-^*mice taking less time to enter the center (Fig. 7). However, the cumulative time spent in the center of the open field was not significant between groups, nor were the elevated plus maze and resident intruder test metrics. Only the difference in attack latency between initial testing and re-testing one day later was significantly different between genotypes. Both genotypes showed a significant preference for the social versus non-social cylinder in the three-chamber task (*p* < 0.0001) but there was no significant difference between the genotypes. Based on this, it seems that a reduction of carnosine content in the mouse OB (∼70%) and brain (∼40%), may result in the *Gadl1^-/-^*mice showing increased initiation to enter an exposed area (which is indicative of decreased anxiety). However, this should be tempered by the observation that the total time spent in the center of the open nor differences in the ratio of the time spent on closed versus open arms of the elevated plus-maze were not different. Further investigation of any possible anxiolytic effect of *Gadl1^-/-^*mice is prudent. To examine the effects of *Gadl1* deletion on organ aging and morphology, we compared tissue sections from 33 mice aged 28-96 weeks, equally matched across different sex and genotypes. At the microscopic level, SKM, brain, and OB morphology were not affected by the elimination of *Gadl1* (Supplementary Fig. 7).

**Fig. 7.**
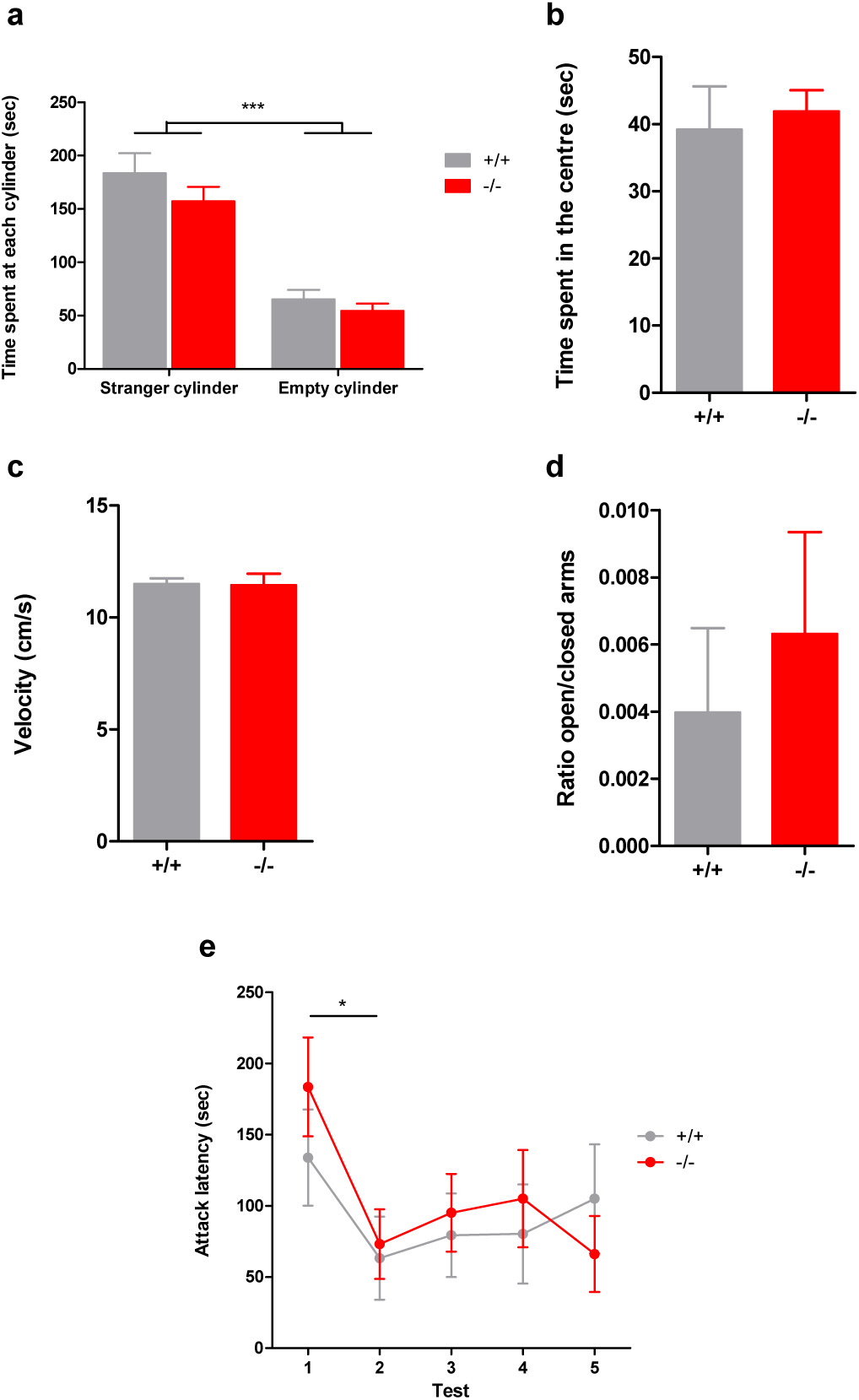
Behavioral phenotypes associated with *Gadl1^-/-^*. **a** 3-chamber task: time spent at each cylinder was not different between the genotypes. Both *Gadl1^+/+^* and *Gadl1^-/-^*mice prefer the social cylinder indicating sociability remains similar. **b** Open Field task: the cumulative time spent in the center was not different between the genotypes. On this measure, no anti-anxiety effect was observed. However, on the latency to enter the center (another anxiety measure), *Gadl1^-/-^*were quicker to enter the center which may suggest some anxiolytic effects of the *Gadl1^-/­^*phenotype which would require confirmation in additional studies. **c** Open Field task: the exploration velocity was not different between the genotypes indicating that no effect on motor function. Similar observations were made with the total distance moved data. Taken together, this suggests no effect of genotype on activity metrics. **d** Elevated plus maze: the ratio of the time spent (s) on the open and closed arms was found not to be different between the genotypes. Both genotypes prefer the closed (sheltered arm) suggesting no difference in anxiety on this measure. **e** Resident Intruder Paradigm: the attack latency against an intruder from the 1^st^ to the 5^th^ day (Test 1-5) was not different between the genotypes. Both genotypes attack faster on the second day compared to the first day after which the attack latency remains constant. This suggests no effect of genotype on aggression.

### Common genetic variants in the *GADL1* locus are associated with multiple human phenotypes

GWA studies have shown associations of the *GADL1* locus with multiple human traits and diseases, including, eating disorders(38), kidney functions(39), and several blood metabolites. This includes a strong association with levels of acetylcarnosine (*p*=8.17e-21) in the *GADL1* locus at an intergenic SNP rs6804368 (13.9 kb from *GADL1* and 17.9 kb from *TGFBR2*)(27). The strongest association (*p*=5.50e-37) has been reported for the response to lithium (Li^+^) in bipolar disorder at an SNP (rs17026688) located in intron 6 of *GADL1*(*23*). However, these findings were not replicated in other clinical samples(25).

To get an overview of genetic associations between human phenotypes and enzymes in carnosine metabolism, we conducted gene-based tests using the MAGMA software(40), mainly focusing on traits related to oxidative protection and brain functions. We included associations of common variants in *GADL1* and seven related genes (Fig. 8)(41). We limited the search to 19 phenotypes with openly available GWAS summary statistics obtained from large samples (total number of tests 151; Bonferroni-corrected *p*-value threshold=3.31e-4). Supplementary Table 3 shows an overview of the data used.

**Fig. 8.**
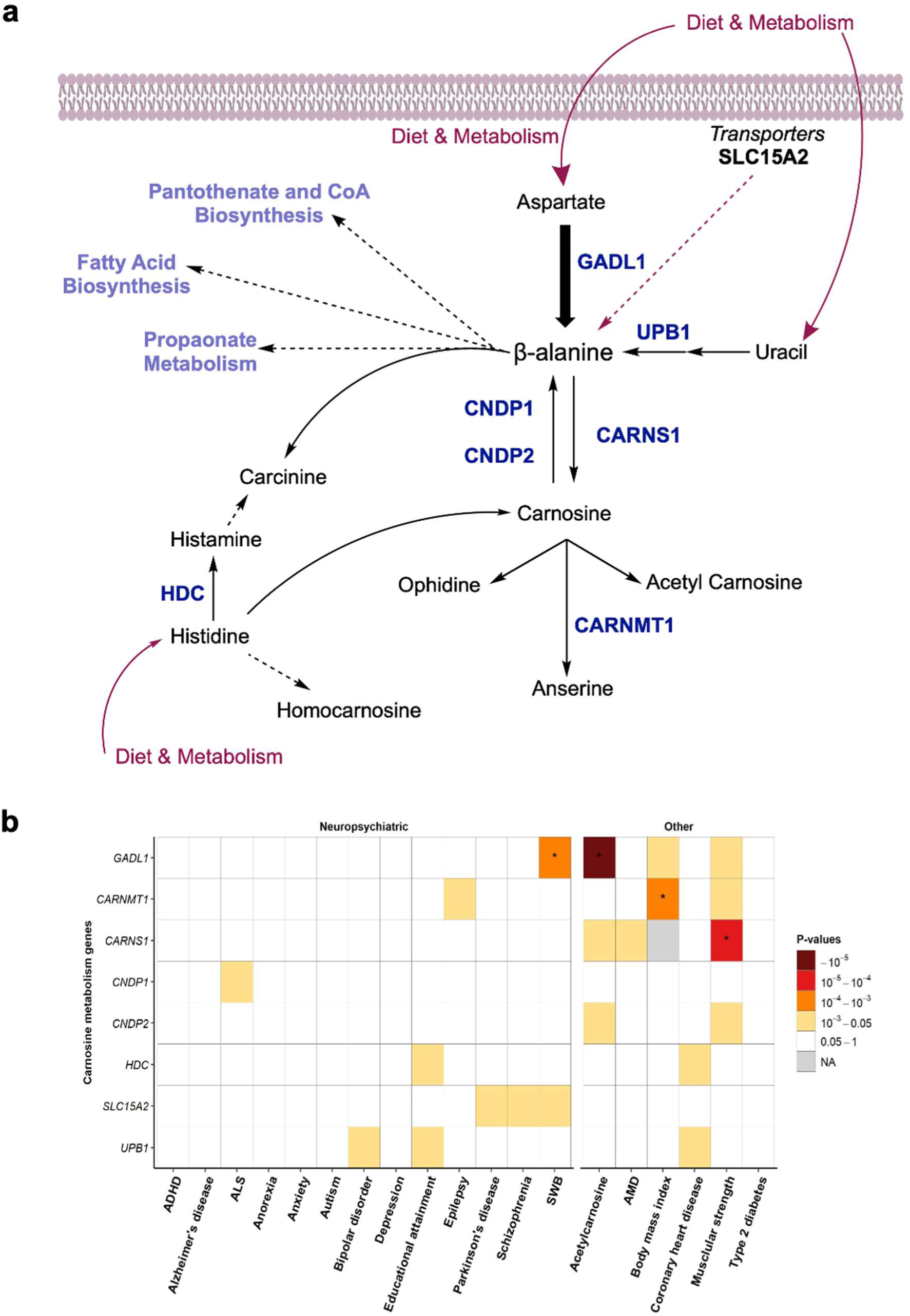
Carnosine homeostasis and human phenotypes. **a** A summary overview of different pathways involving β-alanine. Genes analyzed in this study are marked in blue. **b** Association analyses of genes involved in the carnosine metabolism.

*GADL1* was associated with serum acetylcarnosine levels (*p*=1.25e-11) as well as subjective well-being (SWB) (*p*=1.22e-4). *CARNMT1* was associated with body mass index (BMI), (*p*=1.12e-4). *CARNS1* was associated with right-hand grip strength (muscular strength) (*p*=4.98e-5). In addition, several tentative gene-phenotype associations did not pass the Bonferroni-corrected threshold for significance (Fig. 8b). In conclusion, out of eight tested genes known to be involved in carnosine metabolism, *GADL1* showed the strongest association with human phenotypes. The most obvious associations were observed N-acetylcarnosine levels, muscle strength, and general well-being, but not in a range of neuropsychiatric diseases that have been reported to respond to carnosine treatment^15^.

### DISCUSSION

Carnosine peptides were discovered in 1900, yet their biological functions and biosynthetic routes are still being debated. Here we report the first animal model lacking GADL1. We show that these animals have low levels of β-alanine and carnosine peptides, consistent with a key role of GADL1 in their synthesis. This enzyme is analogous to Asp decarboxylases previously only found in prokaryotes and insects(21).

Although the levels of β-alanine and carnosine peptides were decreased in all tissues examined, this reduction varied from 11% in kidneys to 70-80% in OB tissue. As GADL1 activity was eliminated in *Gadl1^-/-^*mice, the remaining β-alanine and its peptide derivatives may be derived either from the diet, by gut bacteria, or by *de novo* synthesis. Biochemical analyses of the standard plant-based rodent diet did confirm low levels of β-alanine and carnosine, but we cannot exclude a possible small dietary contribution (Supplementary Table 4). It is, however, more likely that there are several alternative biosynthetic routes for β-alanine and that these have different roles in different tissues. Thus, in tissues with the highest demands for carnosine synthesis, i.e. SKM, brain and particularly the OB, we demonstrate a critical role of *de novo* β-alanine biosynthesis to maintain its levels, as well as compensatory upregulation of enzymes involved in alternative biosynthetic routes. These routes may include β-alanine synthesis by CSAD, which has a relatively low β-alanine synthetic specific activity, but much higher tissue abundance than GADL1(25), production of β-alanine from uracil by β-alanine synthase (β-ureidopropionase 1; N-carbamoyl-β-alanine amidohydrolase; (BUP1/UPB1))(42), or by other enzymes, such as β-alanine-2-oxoglutarate transaminase (ABAT) or β-alanine pyruvate transaminase (AGXT2)(22). Our findings contrast earlier observations that found no decarboxylation of Asp in rat muscle(21). Our observation of increased *Upb1* transcript levels in OB tissue of *Gadl1^-/-^* mice also contradicts earlier observations of UPB1 protein in rat liver and kidney, but not in the brain, lung, SKM or spleen(42). Finally, the importance of GADL1 in carnosine synthesis is supported by human genetic studies showing a strong association of *GADL1* variants with blood levels of carnosine peptides(27).

In contrast to the reduced levels of β-alanine, carnosine, anserine, and N-acetylcarnosine, the levels of homocarnosine were increased. This is probably the result of a shortage of β-alanine precursor and carnosine peptides, indicating that the tissue levels of these dipeptides may be tightly regulated. Moreover, no significant difference in GABA between *Gadl1^+/+^* and *Gadl1^-/­^*mice was detected. Together with the observations of increased homocarnosine (a GABA derived peptide), this indicates that GADL1, despite its name, does not catalyze the decarboxylation of glutamate to GABA. Fig. 8a summarizes the different biosynthetic routes of carnosine peptides and the main enzymes and transporters believed to be involved in this metabolism in mammalian tissues.

As the *Gadl1* locus on chromosome 3p24.1-3p23 is close to the *Tgfbr2* locus (3p24.1), and genetic variants in the vicinity of *Gadl1* influence plasma levels of transforming growth factor-beta (TGF-β) receptor type 2 (TGFBR2), the gene targeting strategy was designed to avoid regulatory sequences in this region. This strategy resulted in the expression of low amounts of a catalytically inactive and unstable, partially truncated version of the GADL1 protein, but *Tgfbr2* transcripts were unaltered.

Dietary PLP supplementation significantly increases the concentrations of β-alanine and carnosine peptides in the SKM of rats(22). As GADL1 is dependent on PLP, this increase may be due to the activation of GADL1 by PLP. In contrast, PLP is not required for the production of β-alanine by UPB1. Multiple biosynthetic routes may explain why the elimination of GADL1 only had a modest effect on carnosine peptide levels in SKM (Fig. 2).

The ability of human and mouse GADL1 to catalyze the conversion of Asp to β-alanine was first discovered using recombinantly expressed enzymes with screening against possible physiological substrates(21, 25). However, since the catalytic efficiency of GADL1 *in vitr*o is very low(25), the relevance of this activity *in vivo* has previously been unclear. Here we show that although CSA is the preferred substrate of GADL1 *in vitro*, its decarboxylation of Asp and role in carnosine peptide production is the most striking function in an intact organism.

To determine the substrate specificity of GADL1, we previously expressed the mouse enzyme in bacteria and screened purified GADL1 against all proteinogenic amino acids, as well as many other putative substrates and inhibitors, demonstrating decarboxylase activity against Asp, CSA, and cysteine(25). This substrate profile was similar to what Liu *et al.* observed for the human enzyme(21). However, compared to other PLP-dependent decarboxylases, the affinity and selectivity was very low for all substrates tested. Liu *et al*. reported that it was impossible to detect GADL1 enzyme activities in tissue lysates(21). Here we show that, despite its extremely low activity *in vitro*, GADL1 is important for β-alanine and carnosine peptide synthesis, particularly in the OB, cerebral cortex and SKM. It appears that organs with a high demand for carnosine peptides depend on local synthesis to maintain these levels. *Gadl1^-/­^*mice also have decreased levels of taurine and multiple taurine derivatives in SKM, indicating that CSA also is a physiologically relevant substrate. *Gadl1^-/-^* mice also showed other biochemical alterations which could be secondary to the depletion of β-alanine derivatives and antioxidant function, or primary effects of GADL1 inactivation. *Gadl1* deletion might alter energy metabolism through several pathways. In addition to its role in carnosine synthesis, β-alanine is a component in pantothenic acid (vitamin B5) and coenzyme A, an essential cofactor for multiple biochemical pathways, including energy metabolism.

The role of GADL1 as a relatively non-specific decarboxylase of low molecular weight acid substrates is consistent with its relatively high Km values and low catalytic efficacy(25). Mammalian genomes encode hundreds of PLP-dependent enzymes, many of which may catalyze new or unclassified reactions(32). In addition, due to their common mechanistic features and similar structures, many PLP-dependent enzymes can catalyze multiple biochemical reactions. This promiscuity makes it difficult to define their primary biological functions(32). Some enzymes have evolved to have multiple physiological substrates. Such a reactivity with multiple substrates may be advantageous for fitness(43). GADL1 may be an example of an enzyme with multiple biological activities. Although the strongest effect of GADL1 deletion seemed to be a decrease of β-alanine and carnosine peptides, with the taurine pathway being intact, the *Gadl1^-/-^*mice also had slightly reduced levels of taurine derivatives in SKM, consistent with multiple catalytic functions of this enzyme. Similarly, while the accumulation of oxidative stress biomarkers in *Gadl1^-/-^* mice are probably secondary to loss of β-alanine or carnosine peptides, we cannot exclude that the metabolic changes could be caused by other functions of GADL1.

Inspection of the catalytic site of mouse GADL1 shows that it might be accessible by different small, acidic, polar substrates. From the comparison (Fig. 7), it can be observed that very minor changes in specificity-determining amino acids in the active site vicinity may dictate the function of an enzyme at the tissue and organism level. For a more detailed understanding of the catalytic properties of GADL1, as well as its closest homologues, high-resolution structural studies with active-site ligands, preferably with the flexible catalytic loop visible, will be required.

Both *in vitro* and *in vivo* experiments have shown that reactive oxygen species (ROS) can decrease the levels of antioxidant enzymes, such as superoxide dismutase (SOD) and glutathione peroxidase (GPX), and that carnosine supplementation can restore depleted levels of these enzymes^2^. Carnosine peptides are considered to be important in protection e.g. against ischemia related free radical damage(44). However, the protective function of carnosine has only been studied *in vitro,* or in animals receiving large pharmacological doses of carnosine. Thus, β-alanine and carnosine dietary supplementation have widespread human use and are marketed to protect against oxidative stress(45), (46). However, contradictory findings have been reported, and elevated levels of β-alanine were noted to reduce cellular taurine levels and to be associated with increased oxidative stress(47) and altered energy metabolism(48). Here we show that the *Gadl1* deletion mice have increased levels of oxidative stress biomarkers and altered levels of several antioxidant enzymes. These findings establish this mouse model as a new tool to study not only β-alanine synthesis but also carnosine peptide biology and their relationship to oxidative stress and diseases.

Deletion of the mouse carnosine transporting dipeptide transporter *Pept2 (SLC15A)* has previously been reported to alter carnosine levels in several organs, but not as dramatically as in the *Gadl1^-/-^*mice(49). These organ-specific effects indicate that although β-alanine and carnosine peptides are mainly synthesized locally in the OB, other organs mainly take up carnosine peptides and their precursors from the circulation. Carnosine homeostasis in SKM is regulated by multiple nutritional and hormonal stimuli in a complex interplay between transporters and enzymes, including *CARNS, CNDP2, PHT1, PHT2, TauT, PAT1, ABAT* and *HDC*(*41*) (Fig 7). It was recently reported that in SKM, *CARNS1* and *GADL1* are rapidly upregulated upon high-intensity exercise, demonstrating the importance of carnosine in muscle function and physiological regulation of these enzymes(50).

Since *Gadl1^-/-^*mice displayed behavioral changes and β-alanine and carnosine peptides have been implicated in many different physiological functions and disease states, we performed a gene-based analysis of *GADL1* and enzymes and transporters directly involved in carnosine metabolism. We selected 19 human diseases and traits related to oxidative stress and where treatment with carnosine peptides have shown some promising effects(13) (Fig. 8). In addition to the strong associations with plasma carnosine peptide levels, common variants in the *GADL1* locus were associated with subjective well-being and muscle strength. The association with subjective well-being is particularly intriguing. This phenotype is genetically related to somatic complaints as bodily aches and pains, but also low energy, anxiety and depression, traits that have all been shown in animal studies to be related to levels of carnosine peptides in the brain and SKM(51). The association of *GADL1* with subjective well-being is in line with a carnosine supplementation study(52). However, for most of the diseases studied we observed no significant associations with genes associated with carnosine homeostasis. Still, we cannot exclude possible effects of rare genetic variants or a role in other patient groups or populations. The behavioral data in the *Gadl1^-/-^*mice suggest that the link between *GADL1* with anxiety is worth exploring further but should be interpreted cautiously. Further investigations examining any difference in sensory processing of the environment and alterations in fear-related behavior would be useful next steps.

Brain metabolomics has revealed large regional differences, with particularly large concentration gradients for carnosine(53). However, it is still a mystery why GADL1 and its carnosine peptide products are so abundant in OB. In zebrafish, high immunoreactivity for carnosine and anserine is found in sensory neurons and non-sensory cells of the olfactory epithelium, olfactory nerve, and the OB, indicating a specific role of these peptides in sensory organs(54). Based on the early occurrence of carnosine peptides during embryonic development, it has been suggested that these peptides play a role in the developing nervous system, specifically in the olfactory and visual function(54). Our findings of intact histological architecture and the modest behavioral effects of *Gadl1* deletion may argue against this hypothesis. However, it also possible that the remaining 10-30% carnosine peptides is adequate for maintaining such biological effects, as reported for other neurotrophic factors, such as serotonin(55). Alternatively, a neuroprotective, antioxidant role of GADL1 and carnosine peptides is conceivable. The olfactory epithelium provides a direct entry route from the environment to the OB and brain for substances that could cause oxidative stress. Such substances can cause age-related cellular degeneration and olfactory damage and prevention of such damage depend on an intact antioxidant defense system. Although baseline GADL1 levels are low, the levels of this enzyme may be dynamically regulated in response to oxidative damage and other physiological stressors, as shown for SKM(50).

## Materials

Chemicals in this study were purchased from Sigma Aldrich if not otherwise stated. Probes for the qRT-PCR were from Thermo Fisher, the kits used were from Thermo Fisher or Qiagen, Agarose from Apollo, SDS-Page gels from Bio-Rad. O-Phthaldialdehyde reagent and 2-mercaptoethanol were from Merck–Schuchardt. L-Carnosine nitrate salt was used as standard for NMR spectroscopy.

## Methods

### Ethical statement

This study was carried out per Norwegian laws and regulations, and The European Convention for the Protection of Vertebrate Animals used for Experimental and Other Scientific Purposes. The protocol was approved by the Animal Studies Committee, Norwegian Food Safety Authority (Mattilsynet, permit number 9468). The authors declare no competing interests.

### Generation of *Gadl1*-null mice

*Gadl1*-null (-) mice were generated by homologous recombination in C57BL/6N mice at Genoway, Lyon, France. Based on the *Gadl1* cDNA sequence NM_028638, the exon/intron organization of the gene was established. The mouse *Gadl1* gene is located on chromosome 9 and extends over 183.6 kilo base pairs (kbp) (Sequence Map Chr9:115909455-116074347 bp + strand). The mouse *Gadl1* locus consists of 20 exons (Fig. 1). ATG initiation codons are located in exons 3 and stop codons are located in exons 19 and 20. Bioinformatic analysis identified three isoforms for GADL1 protein and three predicted other splice variants of the *Gadl1 gene*. In the NCBI database release 106, four isoforms, with predicted 518, 479, 362 and 502 amino acid residues (NM_028638) were identified. Based on sequence comparison and X-ray structural data(26), Lys405, Lys333, and His219 are predicted to be essential amino acids of the PLP-binding region and Arg494 and Gln120 are essential amino acids of the substrate-binding region. Exon 7 is highly conserved and contains the substrate-binding residue Gln120. This exon is present in all splice variants. In the mouse genome, the closest coding gene is *Tgfbr2*, encoding the transforming growth factor beta receptor II (Sequence Map Chr9:116087695-116175363 bp, -strand). This is involved in multiple biological pathways and cells. In addition, several non-coding genes and pseudogenes are located in this region. To minimize the risk of interfering with possible regulatory sequences, it was decided to delete only exon 7 in the *Gadl1* gene, resulting in the deletion of the coding sequences encoding the part of the active site domain of *Gadl1*, including Q120(25). The mouse model was generated by homologous recombination in embryonic stem (ES) cells. For this purpose, a targeting vector containing regions homologous to the genomic Gadl1 sequences was constructed. After its transfection into C57BL/6N ES cells, the recombined cell clones were injected into blastocysts. Mouse breeding was established with C57BL/6N Flp-deleter mice to remove the Neomycin cassette. The heterozygous KO colony was produced by breeding with C57BL/6N Cre-deleter mice.

### Breeding

Heterozygous siblings were mated to produce *Gadl1^-/-^*homozygous pups. Further breeding and genotyping were performed at the animal facility at the University of Bergen (Bergen, Norway). Animals were group-housed in temperature-and light-controlled vivarium (21±1 °C; 12/12 h light/dark artificial circadian rhythm). All experiments were conducted with 3 to 96-week-old mice of both sexes that were ad libitum-fed using standard rodent diets (rat and mouse No. 1; RM1) during maintenance and RM3 during breeding (Scanbur/Special Diets Services, Witham, UK). These diets are plant-based. The content of natural amino acids are specified, but the content of carnosine peptides is not specified (http://www.sdsdiets.com/pdfs/RM1-A-P.pdf). No additional analyses of this food were performed.

### *Gadl1* mouse genotype determination

DNA was extracted from mice ear samples at 2 weeks of age. Genotyping was done using a multiplex PCR kit (Qiagen, #206143). After an initial heat-activation at 94 °C for 15 min, DNA was denatured at 94 °C for 30 s, annealed at 60 °C for 1.30 min and extended at 72 °C for 1.30 min. This cycle was repeated 25 times. DNA was extended for an additional 30 min after the last cycle. PCR products were analyzed using 2% agarose gel electrophoresis. The PCR product size was measured to 166 base pairs (bp) in the *Gadl1^-/-^*mice and 330 and 750 bp in the *Gadl1^+/+^* mice. All three bp sizes were measured in the *Gadl1^+/-^*mice (166, 330, and 750 bp). The sizes of PCR products and the primer sequences are summarized in Supplementary Table 5.

### Validation of the animal model

The deletion strategy was validated using DNA sequencing, RNA sequencing, proteomic analyses, and western blotting. DNA sequencing confirmed that exon 7 was absent from the genomic DNA of the *Gadl1^-/-^*mice. We determined branching scores that reflected each exon’s splicing capability using the RNA splicing prediction tool Sroogle (http://sroogle.tau.ac.il/). This gave a lower splicing score for exon 8 compared to exon 9, suggesting that splicing of exon 6 to exon 9 was favored over splicing to exon 8. The qRT-PCR data also suggested that the *Gadl1* mRNA from *Gadl1^-/-^*mice was stable. However, sequence analysis showed that both exon 7 and exon 8 were absent from *Gadl1* mRNA in *Gadl1^-/-^*mice. An mRNA devoid of exons 7 and 8 is expected to be stable and produce a truncated protein. Expression and purification of mouse GADL1 protein in *E. coli* were performed as described(25) (Supplementary Fig. 2). Site-directed mutagenesis of mouse GADL1 to generate a protein lacking exon 7 and 8 (coding protein sequence: NHPRFFNQLYAGLDYYSLAARIITEALNPSIYTYEVSPVFLLVEEAVLKKMIECVGWKEGDGIF NP) was performed using the following primers:

GADL1-R 5’-TTCAGAACCGCCTCTTCCAC-3’

GADL1-F 5’-TCCAAGATTTTTCAACCAGC-3’

GADL1-F 5’-AGAAAGCACCGCCGGCTCCT-3’

GADL1-F 5’-GGATGAGATAGACAGCCTGG-3’

GADL1-F 5’-TTTCTGTTCATGGGGGTAAT-3’

GADL1-R 5’-TGCGGTATTCGGAATCTTGC-3’

GADL1-R 5’-ACGCATCGTGGCCGGCATCA-3’

GADL1-F 5’-CGATTTCGGCCTATTGGTTA-3’

GADL1-R 5’-TGCACAATCTTCTCGCGCAA-3’

GADL1-F 5’-ATGGGGGATCATGTAACTCG-3’

GADL1-R 5’-CTTGCTGCAACTCTCTCAGG-3’

GADL1-F 5’-CGGATCAAGAGCTACCAACT-3’

GADL1-R 5’-TAACGAAGCGCTGGCATTGA-3’

GADL1-F 5’-GCCTTTGAGTGAGCTGATAC-3’

### Metabolomic studies using LC-MS

Animals were anesthetized using 250 mg/mL Urethane administered by intraperitoneal injection and adjusted to their individual weight (1.4-1.8 mg/kg). Tissues were perfused with 50 mL of 0.02% heparin-saline solution pumped through the heart left ventricle before organ extraction. After dissection, the tissues were flash-frozen in liquid nitrogen and stored at −80 °C until use. A total of 172 different tissue samples from 41 animals were shipped at −80 °C to Metabolon Inc (Durham, NC, USA) for further processing. Male and female mice were matched according to genotype and age.

**Table 2.**
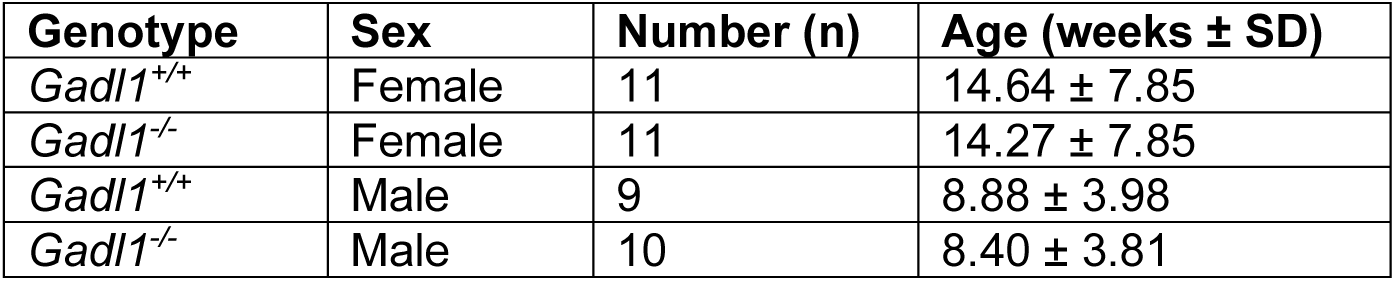
Male and female mice used in the metabolomic study.

All 41 animals were analyzed for brain, SKM, and liver metabolites. A subset of 21 animals (10 *Gadl1^+/+^ and 11 Gadl1^-/-^*; *4.63 and 4.89 weeks, respectively*) was analyzed for OB metabolites. Exploratory analyses were performed in the kidney, cerebellum, heart, and serum from 12-week old males (7 *Gadl1^+/+^ and 7 Gadl1^-/-^*). Due to limited capacity of the animal breeding facility, samples were dissected and shipped in three separate batches, with tissues from 10 (5 *Gadl1^+/+^*, *5 Gadl1^-/-^)*, 17 (8 *Gadl1^+/+^, 9 Gadl1^-/-^)*, and 14 (7 *Gadl1^+/+^*, *7 Gadl1^-/-^)*, animals, respectively. All samples were processed using an identical pipeline

At Metabolon company, samples were prepared using the automated MicroLab STAR® system from Hamilton Company. To remove protein, dissociate small molecules bound to protein or trapped in the precipitated protein matrix, and to recover chemically diverse metabolites, proteins were precipitated with methanol under vigorous shaking for 2 min (Glen Mills GenoGrinder 2000) followed by centrifugation. The resulting extract was divided into four fractions for analysis employing different chromatographic methods. Each sample extract was stored overnight under nitrogen before preparation for analysis. after that, the samples were reconstituted in appropriate solvents (according to the chromatographic method) contained a series of standards at fixed concentrations to ensure injection and chromatographic consistency.

Metabolomic analyses were performed through three chromatographic methods employed were based on reversed-phase (RP) (RP)/UPLC-MS/MS chromatography with positive and negative ion mode ESI and one chromatographic method using HILIC/UPLC-MS/MS with negative ion mode ESI. The columns employed were Waters UPLC BEH C18 (2.1×100 mm, 1.7 µm) and Waters UPLC BEH Amide (2.1×150 mm, 1.7 µm)

Metabolomic analyses were performed at ultra-performance liquid chromatography (UPLC) (Waters ACQUITY) coupled to Thermo Scientific Q-Exactive high resolution/accurate mass spectrometer interfaced with a heated electrospray ionization (HESI-II) source and Orbitrap mass analyzer operated at 35,000 mass resolution. The MS analysis alternated between MS and (56)MS^n^ scans using dynamic exclusion. The scan range varied slighted between methods but covered 70-1000 m/z.

Compounds were identified by comparison to library entries of purified standards or recurrent unknown entities. Metabolon maintains a library based on authenticated standards that contain the retention time/index (RI), mass to charge ratio (*m/z)*, and chromatographic data (including MS/MS spectral data) on all molecules present in the library. Furthermore, biochemical identifications are based on three criteria: retention index within a narrow RI window of the proposed identification, accurate mass match to the library +/- 10 ppm, and the MS/MS forward and reverse scores between the experimental data and authentic standards. The MS/MS scores are based on a comparison of the ions present in the experimental spectrum to the ions present in the library spectrum.

Normalized areas of each compound were rescaled to set the median equal to 1. The values obtained for each tissue were used for metabolomic analyses with Metaboanalyst 4.0. Analyses of partial least squares-discriminant analysis (PLS-DA) data were normalized using Pareto-Scaling.

### ^1^H-NMR spectroscopy

Magic angle spinning (MAS) ^1^H NMR spectroscopy was performed using samples (13-20 mg wet weight) of OB from 12-week old male mice of different genotypes *Gadl1^-/-^, Gadl1^+/-^*and *Gadl1^+/+^.* After dissection, the samples were stored at −80 °C before they were placed in 4 mm ZrO2 MAS rotors with 50 *μ*L inserts and analyzed in a 500 MHz Bruker instrument at 4 °C. Water signal decoupling was achieved by pre-saturation pulses. Each spectrum was recorded at a MAS rate of 5 kHz, using 256 transients and a 5 sec delay between each transient. To aid comparison, the maximal signal intensities were normalized to the same level for all samples.

### ^1^H-MRI of living animals

Animals were anesthetized using 5-6% (v/v) sevoflurane (SevoFlo, Zoetis Inc., Kalamazoo, MI, USA) mixed with oxygen (200 ccm/min) for induction and ∼3% for maintenance. During the MRI experiments, the respiration rate was monitored using a pressure sensor (SA Instruments, NY, USA). MRI investigations were done with User Pharmascan 70/16 (7 T) scanner from Bruker (Bruker BioSpin GmbH, Ettlingen, Germany) using a 72 mm inner diameter transmit coil together with a mouse brain 4-element surface coil array for receiving the MR signal. We recorded 13 coronal T2-weighted images (slice thickness 0.5 mm, slice distance 0.05 mm, 20 mm x 20 mm field of view (FOV), matrix size 256 x 256) with fat suppression. The images were recorded with a turbo-RARE sequence (TE = 38 ms, TR = 3200 ms, 4 averages). The FOV covered the whole brain including the OB. The software used with the scanner was Paravision 6.0.1. Analysis of MR images was done with Fiji ImageJ (version 1.46a, National Institute of Health, Bethesda, MD, USA). The OB was manually delineated in each image slice and the number of pixels within each region of interest was translated to volume measurement.

### RNA-purification and qRT-PCR

Total RNA was purified from different tissues from both *Gadl1^+/+^* and *Gadl1^-/-^*mice using the RNeasy purification kit from Qiagen (#74104 and for muscle tissue #74704). cDNA was synthesized in triplicates from this RNA using a High-Capacity RNA-to-cDNA™ Kit (#4387406, Applied Biosystems™). qRT-PCR was performed using the TaqMan gene expression assay (TaqMan Gene Expression Master Mix #4369016, Applied Biosystems™) The TaqMan probes were: Mm00520087_m1 (CSAD), Mm01348767_m1 (GADL1, exon 13-14), Mm01341531_m1 (GADL1, exon 3-4), Mm01341534_m1 (GADL1, exon 6-7), Mm99999915_g1 (GAPDH) and Mm00607939_s1 (β­actin). mRNA expression for all genotypes was normalized against the housekeeping genes GAPDH and β-actin using a variant (2^ΔCt^) of the Livak Method (2^-ΔΔCt^) as described in the Real-Time PCR applications guide from Bio-Rad(56). Values are presented as mean and upper 95% limit on an Ln y-scale.

### RNA sequencing

RNA sequencing was performed using the Truseq stranded mRNA prep kit from Illumina and an Illumina Hiseq 4000 instrument (paired-end, 75 bp x2 run). Raw data were quality controlled (QC) using the Fastqc tool. (Available at: http://www.bioinformatics.babraham.ac.uk/projects/fastqc). Raw reads were aligned to mouse genome M13 using Hisat2(57). Read aligned to coding regions of the genome were counted using “FeatureCounts”(58). Normalization and differential gene expression was performed using Deseq2(59). Transcripts from 27878 genes at an average were detected. Gene ontology enrichment analysis was performed using tools “TopGo” and their dependencies in R environment (version 2.36.0). Biological pathways affected by the *Gadl1* deletion were defined according to Gene Ontology (GO)(28, 29) and the Kyoto encyclopedia of genes and genomes (KEGG)(28, 60). Among the 27878 mouse genes reliably identified, we further analyzed up-or down-regulated genes with a log2 fold change either less than −1 or greater than +1 and *p*-values ≤ 0.05. Heat maps were made using ClustVis(60).

### Western blotting

We used custom made affinity-purified sheep antibodies generated against purified human GADL1 MBP fusion proteins (James Hastie, University of Dundee). All the other antibodies were purchased from Abcam. The primary antibodies were anti-SOD1 ((CuZn), ab13498, 1:2000), anti-SOD2 ((Mn), ab68155, 1:1000), and anti-GLR1/GSR (ab16801, 1:5000) prepared in 3% (w/v) bovine serum albumin (BSA) in 1X Tris-buffered saline (20 mM Trizma Base, 150 mM NaCl) with 0.1% (v/v) Tween (TBST). Frozen tissue samples (OB, SKM, cerebral cortex) were homogenized in radioimmunoprecipitation assay (RIPA) buffer containing a protease inhibitor cocktail. Equal amounts of protein (30 mg) were separated by SDS-PAGE and transferred to a nitrocellulose membrane (Bio-Rad) using a Trans-Blot Turbo Transfer System (Bio-Rad) according to manufacturer’s protocols. Membranes were blocked with 3% BSA in TBST for 2 h at room temperature (RT) before incubation with primary antibodies overnight at 4 °C. The following day, membranes were incubated with the secondary antibody; anti-rabbit and goat anti-sheep horseradish peroxidase (HRP) IgG H+L (ab6721, 1:10000) (Bio-Rad Laboratories, Hercules, CA) and developed with the enhanced chemiluminescence technique using a Western Bright Sirius kit (Advansta #K-12043-D20) on a Gel DocTM XR+ system (Bio-Rad) with Image Lab software (version 6.0.1). All blots were normalized against the total protein load in a *Gadl1^+/+^* mouse.

### Behavioral Tests

There were 26 male mice, 13 mice from each genotype (*Gadl1^+/+^* and *Gadl1^-/-^*) tested for the behavioral experiments. The average age of *Gadl1^+/+^* and *Gadl1^-/-^* were 22.2 and 21.7 weeks, respectively. The order of testing of the mice was randomized for each test with the tasks following the order, three-chamber social interaction task, open field, elevated plus maze and resident intruder task. Exclusively male mice were used to eliminate possible variation caused by the estrous cycle of female mice. Upon entry, all mice were provided with a unique tail number. All mice were housed at the institutional animal facility in an individually ventilated cage (type 2L, Tecniplast S.p.A., Buguggiate, Italy) with an igloo as environmental enrichment and had *ad libitum* access to water and food. Mice were socially isolated before the open field task. The mice were housed under a reversed light/dark cycle (12/12 h) in a ventilated cabinet, Scantainer (Scanbur, Karlslunde, Denmark) with sunset at 7.30 am at a constant temperature of 24 ± 1°C. All experimental procedures were approved by the Committee of Animal Experiments of the Radboud University Medical Center (project number: DEC2016-0094), Nijmegen, Netherlands.

All the behavioral experiments were performed in the dark phase under red-light conditions in the same experimental room. No experiments were performed within the first hour after the light/dark transition. All animals were given 1 month to acclimatize to the animal housing facility before behavioral testing. EthoVision XT9 software (Noldus, Wageningen, Netherlands) was used for tracking the mice and analysis of the videos in the open field and elevated plus-maze experiments with the Observer version 11 software (Noldus, Wageningen, Netherlands) used for the assessment of the social interaction and resident intruder task experiments. All the behavioral tests were recorded using a high-speed (25 frames per second) infrared camera (GigE, Basler AG, Ahrensburg, Germany). MediaRecorder (Noldus, Wageningen, Netherlands) was used to record these movies.

### Open Field

Locomotion activity was quantified in a 55 × 55 × 36 cm activity chamber which was positioned on a flat table with a camera directly above the center of the apparatus. The animals were placed in the center of the field where locomotion activity was then recorded for 5 min. The arena was divided into four quadrants in which the connected center points of all quadrants formed the center of the field and measured 27.5 × 27.5 cm. The total time in the center zone, outside the center and time spent near the walls, was measured as well as the frequency of center visits. In addition, the latency to leave the center was used as an indication of immobility behavior. Reduced frequency of center visits, velocity, and distance traveled was used as indications of locomotor activity and anxiety behavior. Total distance traveled was measured by tracking movement from the center of the mice’s body. The arena was cleaned with 70% alcohol between tests. Videotapes of the locomotion activity were examined using EthoVision XT9 (Noldus, Wageningen, Netherlands).

### Elevated plus-maze

The apparatus provided by Noldus consisted of two open arms and two closed arms (36 cm × 6 cm, walls 15-cm-high) with a common central platform (6 cm^2^). The apparatus was placed 60 cm above the ground. At the start, animals were placed at the junction of the open and closed arms with the head facing the closed arms. The elevated plus-maze was performed in the dark phase under dim-light conditions. The time spent in the open and closed arms were examined as a time ratio (RT). The RT is the time spent in the open arms (TO)/total time spent in both closed (TC) and open arms (TO): RT = TO/(TO + TC). Furthermore, the frequency of transition between the arms, the total distance traveled, and velocity was measured. Between each animal, the maze was cleaned with 70% ethanol and dried before testing the next animal.

### Resident–Intruder Paradigm

The resident–intruder paradigm was performed to assess territorial aggression. On each testing day, an unfamiliar C57BL/6J intruder mouse was encountered, which was randomly assigned to a resident for each interaction. All animals, both resident and intruder, were tested once a day. The housing cage of the resident was used as the interaction area. A transparent Plexiglas screen was placed in the middle of the cage, to prevent direct interaction between animals but to enable visual, auditory, and olfactory perception. The intruder mouse was placed at the other side of the plastic screen for a period of 5 min. Hereafter, the screen was removed, and the interaction was videotaped for 5 min. After the test, the intruder was removed from the cage, and both animals were weighed and checked for wounds. The frequency of attacks and bites and the latency to the first attack were analyzed manually for all interaction days. An attack latency of 300 s was taken in case no attack occurred within the 5-min interaction window.

### Three-Chamber Social Interaction Test

A standard 3 chamber task arena (Noldus, Wageningen, Netherlands) was used in a dark room with red overhead lighting. During the first 10 min of the test, the test mouse was placed alone in the arena to habituate to the new environment. In both the left and right chamber an empty acrylic cylinder with bars was placed. After 10 min, an interaction mouse (C57BL/6J mouse, same age as test mouse) was placed in a cylinder, randomly in either the left or right cylinder. Testing was spread across 2 days with each C57BL/6J mouse used maximally 4 times as an interaction mouse. The order of testing was counterbalanced. For 10 min, test mice were allowed to investigate the arena with the interaction mouse in it.

### Carnosine related genes in human phenotypes

To conduct the gene-based analyses, we obtained openly available summary statistics from GWAS performed in individuals of European descent (Supplementary table 3). We then restricted the data, where possible, to SNPs with minor allele frequency 1%, with good imputation quality (INFO > 0.8) and that were represented in more than 70% of the total meta-analyzed sample size. We then calculated the eight genes of interest’s association with the chosen phenotypes using MAGMA(40). The gene-phenotype associations were calculated using the individual SNPs’ P-values for association with the respective phenotypes. SNPs were assigned to a gene if their chromosomal location were within the start and end of the gene transcripts (i.e. the standard settings of MAGMA). We used the 1000 genomes CEU population as the reference panel to correct for linkage disequilibrium.

### Variant Effect Prediction

In total, Ensembl reported 23,292 SNPs in human GADL1, with 78% of them being intronic. Among the coding variants, 67% were presented as missense, 23% as synonymous, 5% as frameshift, 3% as stop-gained and 1% as in-frame deletion.

### Structural analyses

For comparative protein structure analysis, the structures of mouse GADL1(26) (PDB entry 6enz), human CSAD (PDB entry 2jis), and human GAD65(61) (PDB entries 2okk/2okj) were used. To predict the binding mode of Glu into GAD65, its structure was re-refined and observed to contain the inhibitor chelidonic acid, based on which the Glu binding mode can easily be obtained, due to the presence of two appropriately spaced carboxyl groups. Structures were superposed in COOT(62) and visualized in UCSF Chimera(63).

### Hematoxylin and eosin staining

For the histochemistry studies, the organs were fixed with formalin. The samples were paraffin-embedded and sectioned into 40 mm thick slices. For hematoxylin and eosin (HE) staining, sections were deparaffinized in Tissue-Clear II (2×10 min) and rehydrated using 100% ethanol (2×1 min each), followed by 96% ethanol (1 min) and then 70% ethanol for 1 min and ddH_2_O for 3 min. Sections were then dipped in hematoxylin 0.2% (Histolab) for 5 min, eosin for 1 min, followed by a 5 sec wash with ddH_2_O. The sections were then rehydrated in 70% ethanol for 30 sec, followed by dipping in 96% (2 min) and 100% ethanol (2×2 min). Finally, the sections were dipped in Xylene for 2 min, mounted on slides and coverslipped. Hamamatsu slide scanner was used for scanning slides and Aperio software was used to take pictures.

## Statistics

All data, unless specified otherwise, are presented as mean ± standard error (SEM) or mean ± standard deviation (SD). The growth curves (Fig. 1b-c), enzyme activity assay (Fig. 1h), NMR (Fig. 3c), MRI and animal behavior (Fig. 6) was analyzed with an unpaired Student’s t-test (two-tailed). Statistical significance was defined as p ≤ 0.05. Metabolomics data (Fig.2, Supplementary Fig. 4-6) was analyzed with an unpaired Student’s t-test (two-tailed) with Welch’s correction. Antioxidant levels (Fig. 5) were analyzed with a ratio, paired t-test (two-tailed) per experimental setup. All experiments were run in technical and biological triplicates.

## Study approval

The protocol used in this manuscript, was approved by the Animal Studies Committee, Norwegian Food Safety Authority (Mattilsynet, permit number 9468)

## Acknowledgements

We thank Lars E. Schiro, Hans Olav Rolfsnes, Hege Dale, and Bendik Nordanger for expert technical assistance. This work has received funding from Stiftelsen Kristian Gerhard Jebsen (SKJ-MED-02) and The Regional Health Authority of Western Norway (No. 25048). The Genomics Core Facility (GCF) at the University of Bergen, which is a part of the NorSeq consortium, provided services on RNA sequencing. GCF is supported in part by major grants from the Research Council of Norway (grant no. 245979/F50) and Trond Mohn Stiftelse (grant no. BFS2016-genom).

## Author contribution

E.M. characterized all aspects of the mutated mice, performed mice dissection, genotyping, enzyme and western blotting assays, analyzed the MR and NMR data and wrote the manuscript; S.C.H. performed western blotting assays, analyzed mouse tissue lysates and RNA data and helped with the preparation of the manuscript; R.K. helped in the analyses of metabolomic data; I.W. performed genotyping and qRT-PCR; T.-A.H. performed the gene-based human analyses and helped in the analyses of metabolomic data; R.M.P. performed the statistical analyses of metabolomic data; C.T. performed the NMR spectroscopic experiments; F.M. performed the mouse behavioral studies; A.B. performed the qRT-PCR studies; J.G. supervised the mouse behavioral studies; H.M. performed histological examination of mouse tissues; P.K. analyzed and compared the protein structures; J.H. conceived and supervised the study and wrote the manuscript.

## Conflict of interest

The authors have declared that no conflict of interest exists.

## Data availability

All data generated or analyzed during this study are available from the authors.

