## Supplemental table 1 for "GADL1 is a multifunctional decarboxylase with tissue-specific roles in β-alanine and carnosine production"

### Supplementary tables

**Supplementary Table 1a** Top 25 upregulated genes in OB tissue samples from *Gadl1*$^{-/-}$ compared to *Gadl1*$^{+/+}$ mice.

| ENSEMBL ID | Abbreviation | Name | Log2Fold | P-value |
|---|---|---|---|---|
| ENSMUSG00000032226 | Gcnt3 | Glucosaminyl (N-acetyl) transferase 3, mucin type | 6.05 | 0.00084 |
| ENSMUSG00000020159 | Gabrp | γ-aminobutyric acid (GABA) A receptor, pi | 5.77 | 0.01167 |
| ENSMUSG00000085224 | Gm13425 | Predicted gene 13425 | 5.40 | 0.00262 |
| ENSMUSG00000038805 | Six3 | Sine oculis-related homeobox 3 | 5.23 | 0.00077 |
| ENSMUSG00000093894 | Ighv1-53 | Immunoglobulin heavy variable 1-53 | 5.22 | 0.02869 |
| ENSMUSG00000096225 | Lhx8 | LIM homeobox protein 8 | 5.17 | 0.00055 |
| ENSMUSG00000045620 | Odf3l1 | Outer dense fiber of sperm tails 3-like 1 | 4.97 | 0.00314 |
| ENSMUSG00000061959 | Ces1e | Carboxylesterase 1E | 4.91 | 0.00405 |
| ENSMUSG00000004341 | Gpx6 | Glutathione peroxidase 6 | 4.86 | 0.00372 |
| ENSMUSG00000029866 | Kel | Kell blood group | 4.67 | 0.00028 |
| ENSMUSG00000046975 | Olfr1020 | Olfactory receptor 1020 | 4.57 | 0.01397 |
| ENSMUSG00000027902 | Chil6 | Chitinase like 6 | 4.55 | 0.00405 |
| ENSMUSG00000024681 | Ms4a3 | Membrane-spanning 4-domains A3 | 4.49 | 0.02221 |
| ENSMUSG00000112343 | Sfta3-ps | Surfactant associated 3, pseudogene | 4.38 | 0.02155 |
| ENSMUSG00000082308 | Gm15770 | Predicted gene 15770 | 4.36 | 0.03261 |
| ENSMUSG00000083986 | Gm12213 | Predicted gene 12213 | 4.36 | 0.02563 |
| ENSMUSG00000090475 | Gm6245 | Predicted gene 6245 | 4.36 | 0.02527 |
| ENSMUSG00000074665 | Bpifb4 | BPI fold containing family B4 | 4.35 | 0.01778 |
| ENSMUSG00000073920 | Olfr661 | Olfactory receptor 661 | 4.30 | 0.02962 |
| ENSMUSG00000058884 | Olfr1025-ps1 | Olfactory receptor 1025, pseudogene 1 | 4.30 | 0.03356 |
| ENSMUSG00000087340 | Gm15228 | Predicted gene 15228 | 4.25 | 0.02579 |
| ENSMUSG00000094872 | Igkv9-120 | Immunoglobulin kappa chain variable 9-120 | 4.22 | 0.01533 |
| ENSMUSG00000066108 | Muc5b | Mucin 5, subtype B, tracheobronchial | 4.21 | 0.00077 |
| ENSMUSG00000105906 | Iglc1 | Immunoglobulin lambda constant 1 | 4.18 | 0.01064 |
| ENSMUSG00000095765 | Olfr741 | Olfactory receptor 741 | 4.16 | 0.00863 |

**Supplementary Table 1b** Top 25 downregulated genes in OB tissue samples from *Gadl1*$^{-/-}$ compared to *Gadl1*$^{+/+}$ mice.

| ENSEMBL ID | Abbreviation | Name | Log2Fold | P-value |
|---|---|---|---|---|
| ENSMUSG00000022485 | Hoxc5 | Homeobox Protein Hox C5 | -8.25 | 0.00033 |
| ENSMUSG00000038700 | Hoxb5 | Homeobox Protein Hox B5 | -7.65 | 0.00325 |
| ENSMUSG00000087658 | Hotairm1 | Hoxa transcript antisense RNA, myeloid-specific 1 | -6.90 | 0.00079 |
| ENSMUSG00000038253 | Hoxa5 | Homeobox Protein Hox A5 | -6.46 | 0.00381 |
| ENSMUSG00000056423 | Uts2b | Urotensin-2B | -6.04 | 0.00188 |
| ENSMUSG00000048763 | Hoxb3 | Homeobox Protein Hox B3 | -5.99 | 0.00081 |
| ENSMUSG00000001661 | Hoxc6 | Homeobox Protein Hox C6 | -5.97 | 0.01110 |
| ENSMUSG00000056468 | 5730596B20Rik | RIKEN cDNA 5730596B20 gene | -5.84 | 0.03447 |
| ENSMUSG00000084844 | Hoxb3os | Homeobox B3 and homeobox B2, opposite strand | -5.76 | 0.03083 |
| ENSMUSG00000075394 | Hoxc4 | Homeobox Protein Hox C4 | -5.67 | 0.00327 |
| ENSMUSG00000085696 | Hoxaas3 | Hoxa cluster antisense RNA 3 | -5.02 | 0.00906 |
| ENSMUSG00000060738 | Prl7c1 | Prolactin 7c1 | -4.85 | 0.00353 |
| ENSMUSG00000005503 | Evx1 | Even-skipped homeobox 1 | -4.65 | 0.03972 |
| ENSMUSG00000067684 | Obp1a | Odorant binding protein IA | -4.55 | 0.02233 |
| ENSMUSG00000001670 | Tat | Tyrosine aminotransferase | -4.48 | 0.02410 |
| ENSMUSG00000103430 | Gm36996 | Predicted gene 36996 | -4.45 | 0.01718 |
| ENSMUSG00000108282 | Gm44317 | Predicted gene 44317 | -4.41 | 0.01906 |
| ENSMUSG00000026976 | Pax8 | Paired box 8 | -4.29 | 0.00069 |
| ENSMUSG00000038155 | Gstp2 | Glutathione S-transferase, pi 2 | -4.28 | 0.03740 |
| ENSMUSG00000042279 | H1foo | H1.8 linker histone | -4.27 | 0.02103 |
| ENSMUSG00000109753 | Gm45633 | Predicted gene 45633 | -4.26 | 0.02596 |
| ENSMUSG00000041333 | Mup4 | Major urinary protein 4 | -4.26 | 0.02462 |
| ENSMUSG00000073242 | Dnmt3aos | DNA methyltransferase 3A, opposite strand | -4.18 | 0.02929 |
| ENSMUSG00000074385 | Gm10684 | Predicted gene 10684 | -4.14 | 0.00057 |
| ENSMUSG00000029844 | Hoxa1 | Homeobox Protein Hox A1 | -4.11 | 0.03148 |

**Supplementary Table 3.** The overview of the data used for genetic analysis using MAGMA software.

| Condition | Sample size* | Reference and data source |
|---|---|---|
| Neuropsychiatric | | |
| ADHD** | 53,293 | Demontis et al. [1] |
| Alzheimer's disease | 452,010 | Jansen et al. [2] |
| ALS*** | 80,610 | Nicolas et al. [3] |
| Anorexia | 72,517 | Watson et al. [4] |
| Anxiety | 18,186 | Otowa et al. [5] |
| Autism | 46,351 | Grove et al. [6] |
| Bipolar disorder | 51,710 | Stahl et al. [7] |
| Depression | 173,005 | Wray et al. [8] |
| Educational attainment | 766,345 | Lee et al. [9] |
| Epilepsy | 34,852 | International League Against Epilepsy Consortium on Complex Epilepsies [10] |
| Parkinson's disease | 482,730 | Nalls et al. [11] |
| Schizophrenia | 77,096 | Ripke et al. [12] |
| SWB**** | 298,420 | Okbay et al. [13] |
| Acetylcarnosine (in blood serum) | 6279 | Shin et al. [14] |
| AMD | 33,976 | Fritsche et al.[15] |
| Body mass index | 795,640 | Yengo et al.[16] |
| Coronary heart disease | 547,261 | van der Harst & Verweij [17] |
| Muscular strength****** | 335,842 | UKBIOBANK http://www.nealelab.is/uk-biobank http://ldsc.broadinstitute.org/ |
| Type 2 diabetes | 605,056 | Xue et al. [18] |

*Maximum total sample size

** Attention-deficity/hyperactivity disorder

*** Amytrophic lateral sclerosis

****Subjective well-being

***** Age-related macular degeneration

****** Right hand grip strength

**Supplementary Table 4** Diet analysis.

| Parameter analyzed | Result (g/kg) |
|---|---|
| 1-Methylhistidine (Free) | <0.5 |
| 3-Methylhistidine (Free) | <0.5 |
| Alanine (Free) | <0.5 |
| Alpha-Aminoadipic acid (Free) | <0.5 |
| Alpha-Amino-n-butyric acid (Free) | <0.5 |
| Anserine (Free) | <0.5 |
| Arginine (Free) | <0.5 |
| Asparagine (Free) | 0.535 |
| Aspartic Acid (Free) | <0.5 |
| β-Alanine (Free) | <0.5 |
| β-Aminoisobutyric acid (Free) | <0.5 |
| Carnosine | <0.5 |
| Citrulline | <0.5 |
| Cystathionine (Free) | <0.5 |
| Cystin (Free) | <0.5 |
| Delta-Hydroxylysine (Free) | <0.5 |
| Ethanolamine | <0.5 |
| γ-Amino-butyric acid (Free) | <0.5 |
| Glutamic acid (Free) | <0.5 |
| Glutamine (Free) | <0.5 |
| Glycine (Free) | <0.5 |
| Histidine (Free) | <0.5 |
| Homocysteine (Free) | <0.5 |
| Hydroxyproline (Free) | <0.5 |
| Isoleucine (Free) | <0.5 |
| Leucine (Free) | <0.5 |
| Lysine (Free) | 0.843 |
| Methionine (Free) | <0.5 |
| Ornithine (Free) | <0.5 |
| Phenylalanine (Free) | <0.5 |
| Phosphothanolamine (Free) | <0.5 |
| Phosphoserine (Free) | <0.5 |
| Proline (Free) | <0.5 |
| Sarcosine (Free) | <0.5 |
| Serine (Free) | <0.5 |
| Taurine (Free) | <0.5 |
| Threonine (Free) | <0.5 |
| Tryptophan (Free) | <0.5 |
| Tyrosine (Free) | <0.5 |
| Urea | <0.5 |

**Supplementary Table 5.** The sequence of the PCR primers and the size of the PCR product.

| Primer | Primer Sequence 5'-3' | PCR Product Size | |
|---|---|---|---|
| | | WT | KO |
| 136258Cre-HAA2 | TCAGTTGAGAAGCCCTTCCTTGGTGTA | 330 and 750 bp | 166 bp |
| 136249Cre-HAA2 | CCTTGAACGTGGTTCTCTAGTAGCCACC | | |
| 136248Cre-HAA2 | AGACCTGGTTAAGCAACTCTCCACTAACTCC | | |

#### Supplementary Figures

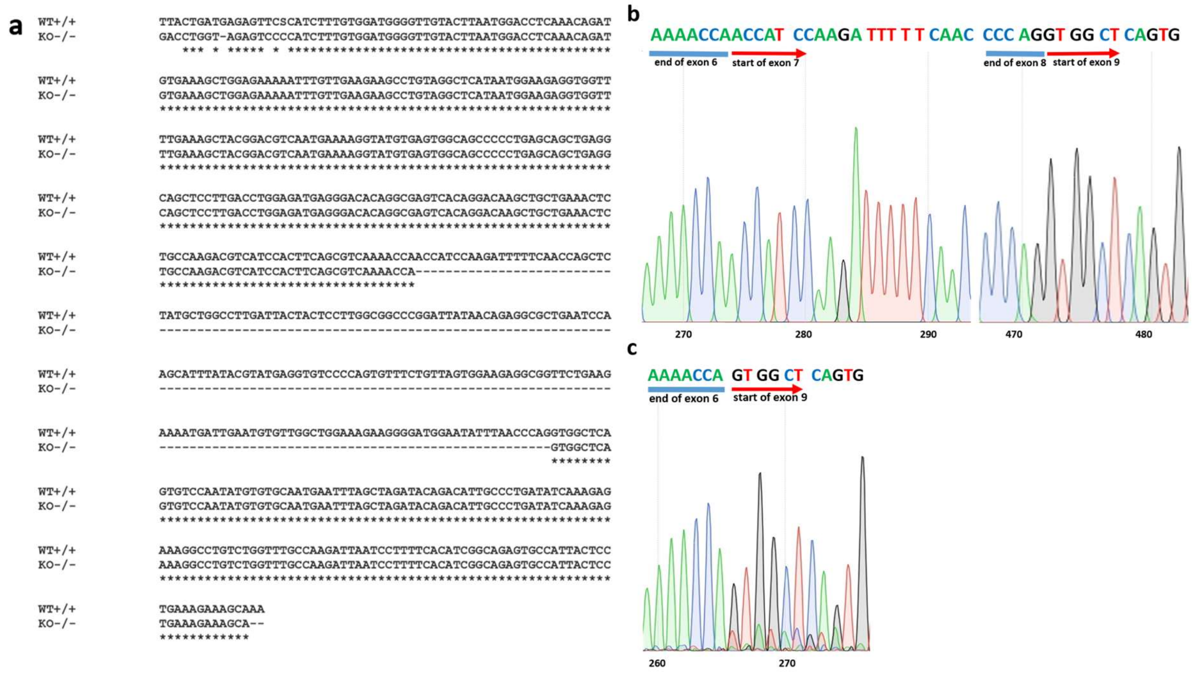


**Supplementary Fig. 1** Sequencing of *Gadl1* mRNA confirming that exon 7 and 8 are missing from the *Gadl1*$^{-/-}$ muscle transcript. **a** Alignment of mRNA sequences of the *Gadl1*$^{+/+}$ and *Gadl1*$^{-/-}$ mice. Cluster omega was used for alignment (https://www.ebi.ac.uk/Tools/msa/muscle/). Created in SnapGene (v4.2, GSL Biotech). **b,c** sequencing chromatogram of **(b)** *Gadl1*$^{+/+}$ and **(c)** *Gadl1*$^{-/-}$ mice. Exon 7 and 8 are missing in the *Gadl1*$^{-/-}$ mice mRNA.

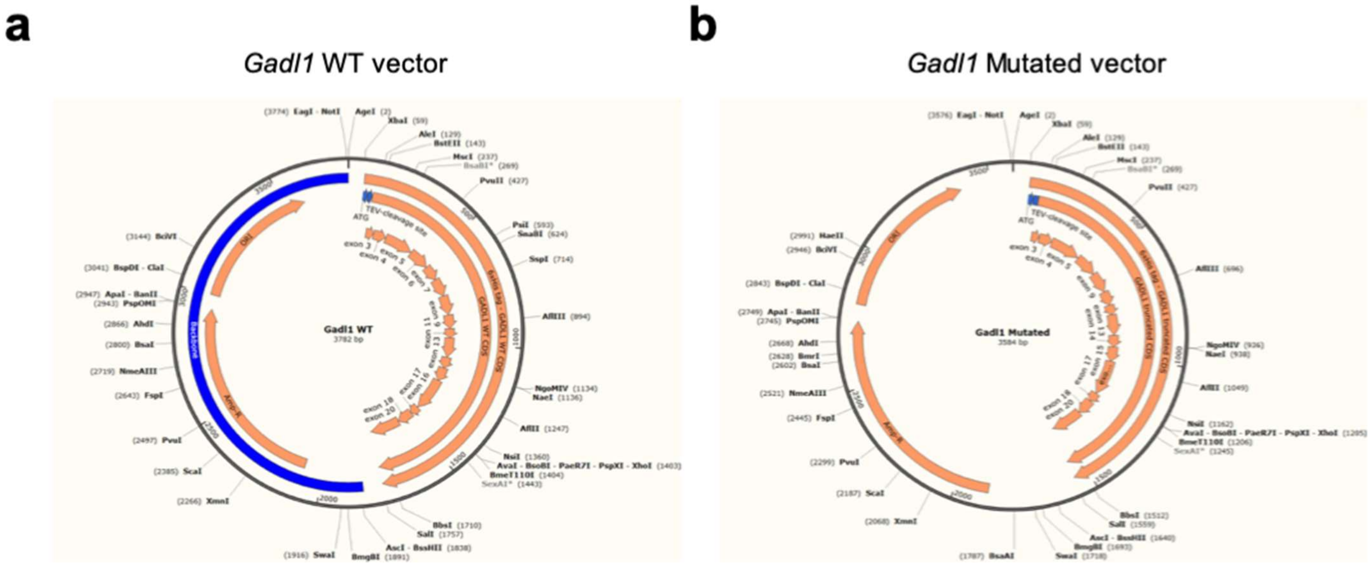


**Supplementary Fig. 2** Vector design. **a-b** *Gadl1* **(a)** wild type (WT) and *Gadl1* **(b)** mutated vector. The vectors were made in BL21 E-coli. The proteins were expressed and purified using Histidine-tag. The purified enzymes were tested with an activity assay. Created in SnapGene (v4.2, GSL Biotech).

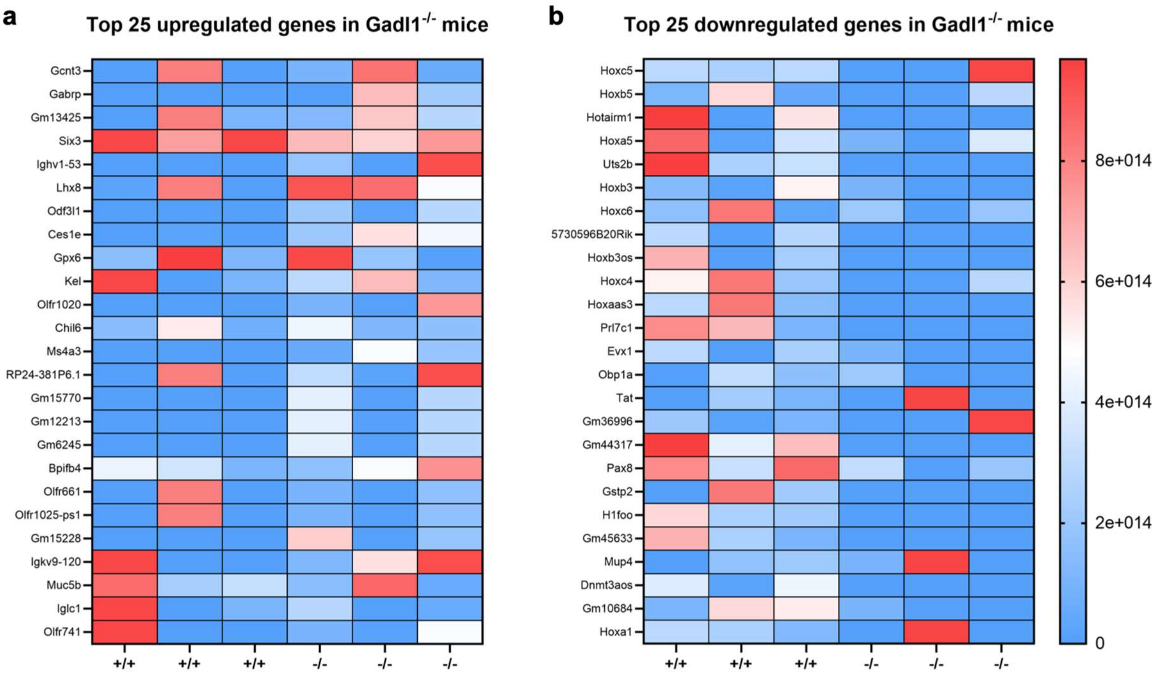


**Supplementary Fig. 3** Top 25 up- and down-regulated genes in olfactory bulb when comparing *Gadl1*$^{-/-}$ (n =3) to *Gadl1*$^{+/+}$ (n = 3).

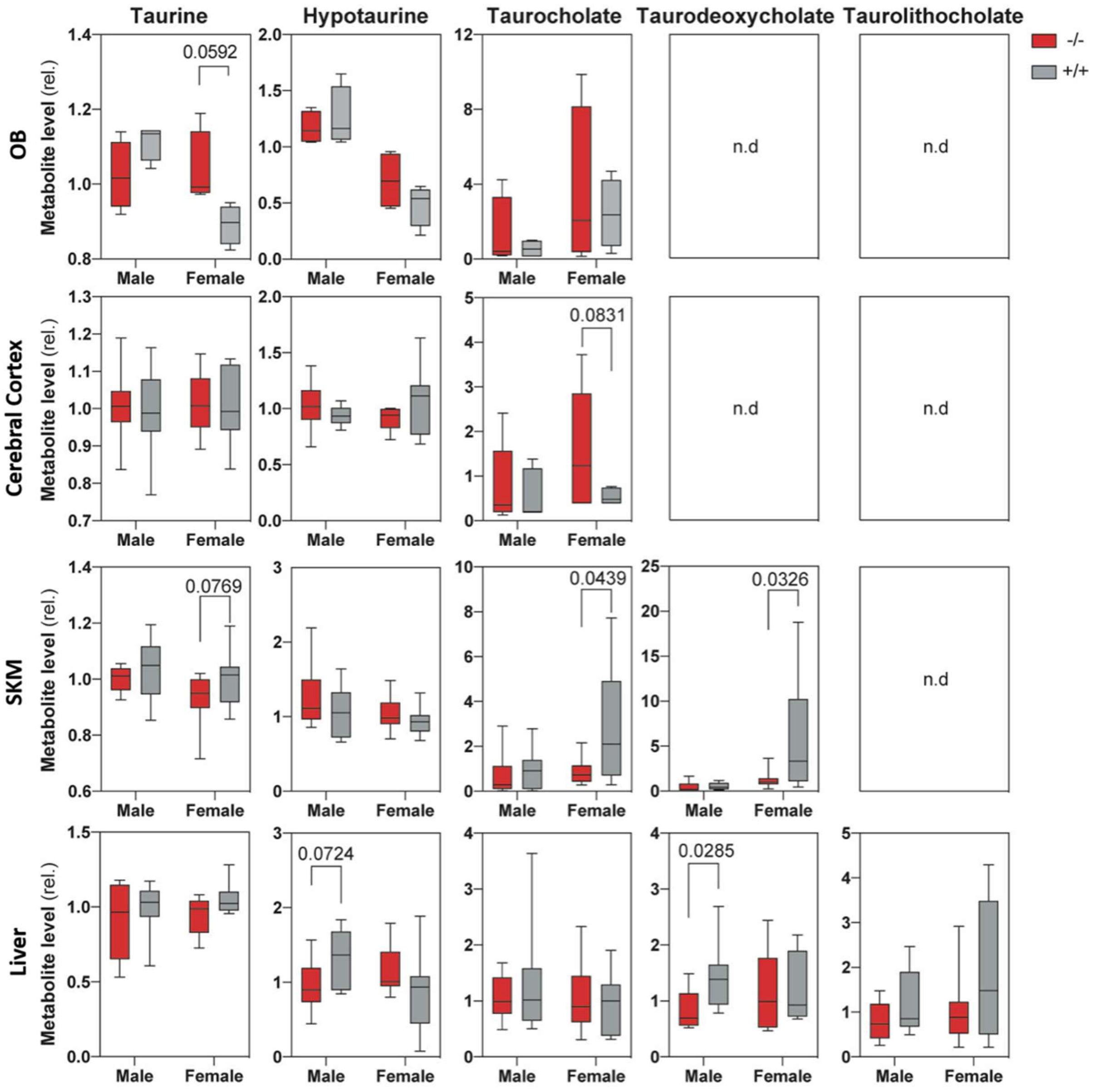


**Supplementary Fig. 4** Distribution of Taurine and taurine derivatives in *Gadl1*$^{+/+}$ and *Gadl1*$^{-/-}$ mice (male and female)

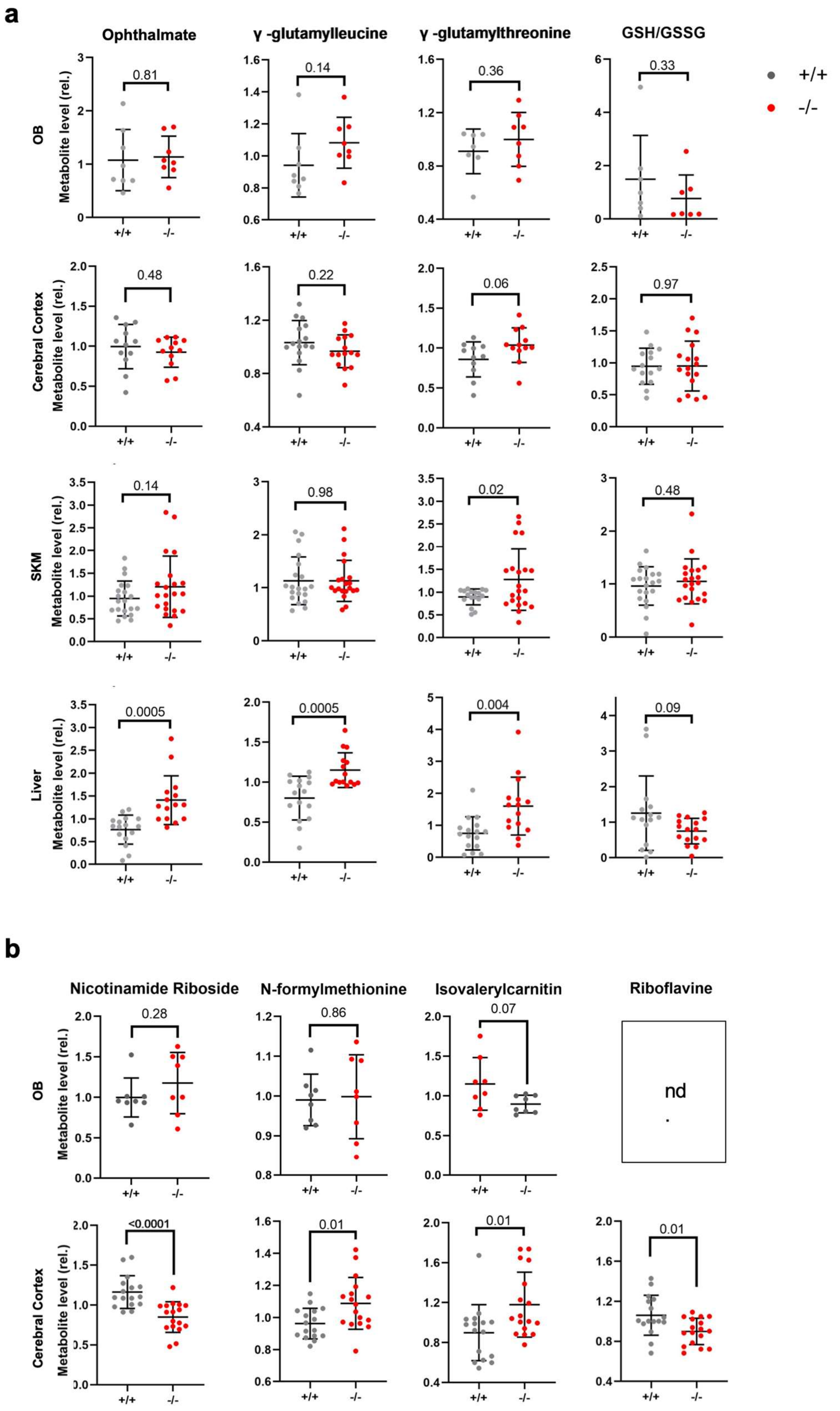


**Supplementary Fig. 5** Glutamine peptides and oxidative stress markers. Untargeted metabolomic profiling of tissue samples from *Gadl1*$^{+/+}$ (grey) and *Gadl1*$^{-/-}$ (red) mice (female and male).

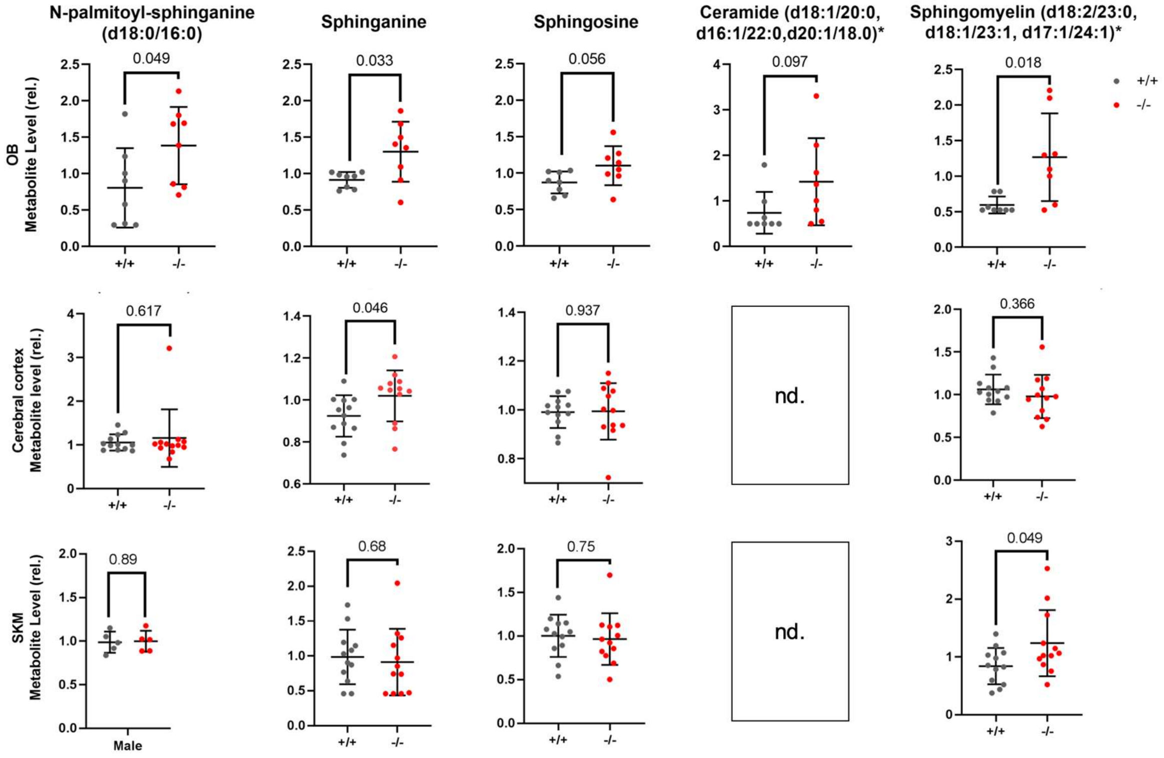


**Supplementary Fig. 6** Effect of Gadl1 deletion on sphingolipid and ceramide metabolism. Untargeted metabolomic profiling of tissue samples from *Gadl1*$^{+/+}$ (grey) and *Gadl1*$^{-/-}$ (red) mice (female and male).

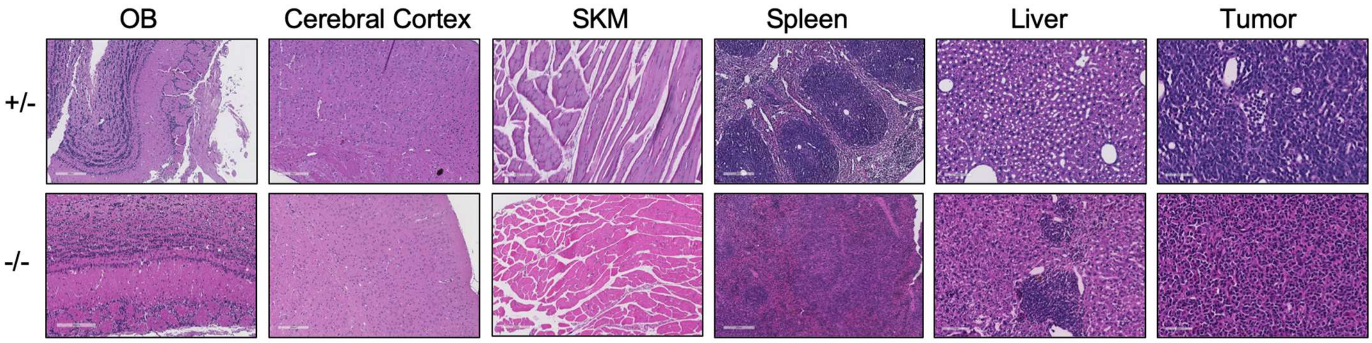


**Supplementary Fig. 7.** HE staining. Representative images of OB, cerebral cortex, SKM, spleen (all 10X, 200 μm), liver (10X, 100 μm), and tumor (40X, 50 μm) morphology in *Gadl1*$^{+/-}$ (male, 74 weeks) and *Gadl1*$^{-/-}$ (female, 96 weeks).
